# Newly born mesenchymal cells disperse through a rapid mechanosensitive migration

**DOI:** 10.1101/2023.01.27.525849

**Authors:** Jon Riddell, Shahzeb Raja Noureen, Luigi Sedda, James D. Glover, William K. W. Ho, Connor A. Bain, Arianna Berbeglia, Helen Brown, Calum Anderson, Yuhang Chen, Michael L. Crichton, Christian A. Yates, Richard L. Mort, Denis J. Headon

## Abstract

Embryonic mesenchymal cells are dispersed within an extracellular matrix but can coalesce to form condensates with key developmental roles. Cells within condensates undergo fate and morphological changes, and induce cell fate changes in nearby epithelia to produce structures including hair follicles, feathers or intestinal villi. Here, by imaging of mouse and chicken embryonic skin, we find that mesenchymal cells undergo much of their dispersal in early interphase, in a stereotyped process of displacement driven by three hours of rapid and persistent migration, followed by a long period of low motility. The cell division plane and the elevated migration speed and persistence of newly born mesenchymal cells are mechanosensitive, aligning with tension in the tissue. This early G1 migratory behaviour disperses mesenchymal cells and allows the daughters of recent divisions to travel long distances to enter dermal condensates, demonstrating an unanticipated effect of a cell cycle sub-phase on core mesenchymal behaviour.

**Highlights:**

- After mesenchymal cell division the speed and persistence of daughter cell migration is elevated for 180 minutes
- Mesenchymal cell division and migration are directed by tissue tension
- Newly born mesenchymal cells are uniquely responsive to tissue strain
- Newly born mesenchymal cells are preferentially recruited to dermal condensates
- Increased dispersal of newly born cells enables long distance travel to dermal condensates

**Graphical abstract:** 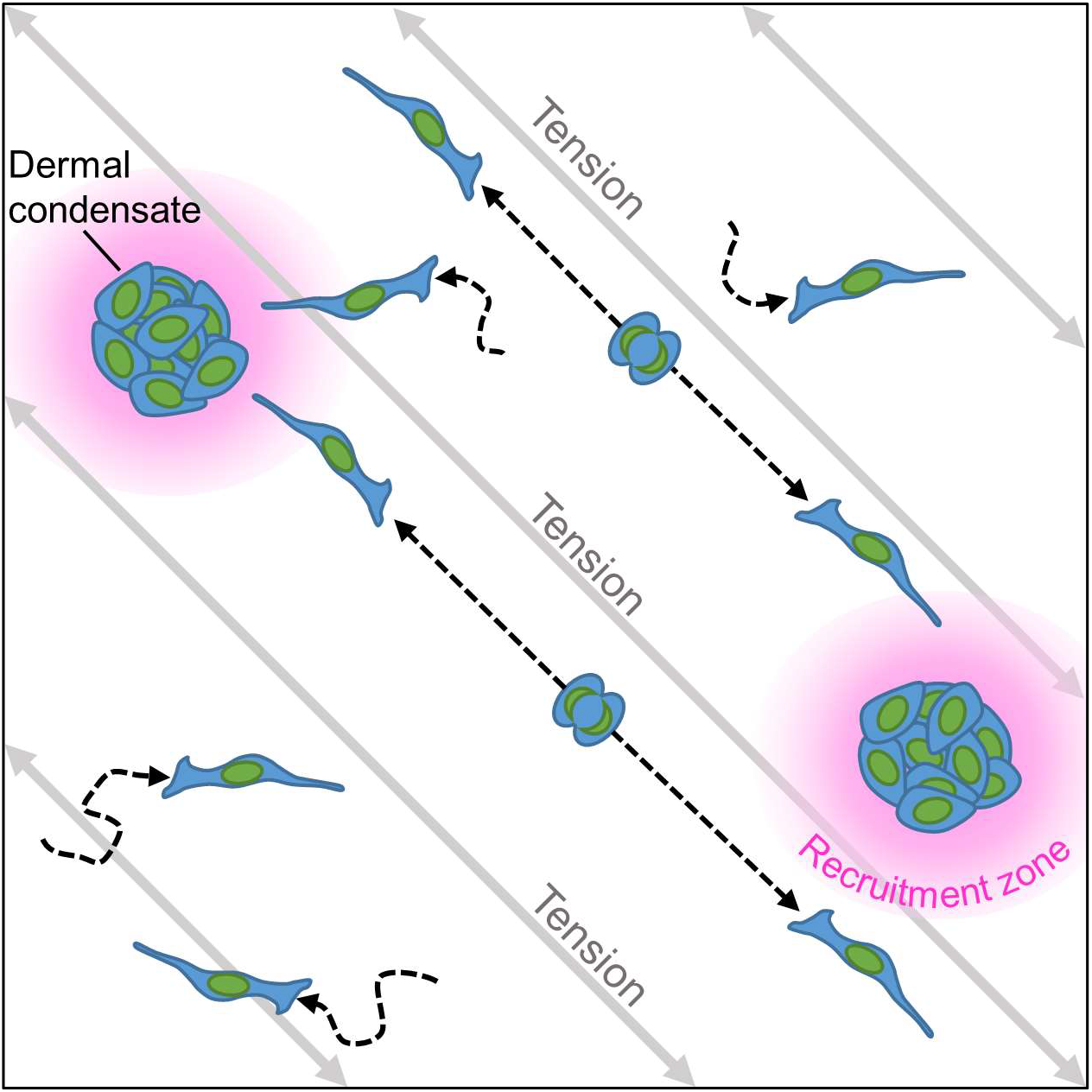

## Introduction

Many vertebrate organs, including lung, intestine, skin, kidney and mammary gland, are composites of an epithelium and a connective tissue stroma. The stromal component originates from embryonic mesenchyme, consisting of motile cells dispersed within an extracellular matrix. In the skin the mesenchyme is largely populated with embryonic fibroblasts, while also carrying adipocytes, blood vessels, lymphatics, nerves, macrophages and melanocytes^1^. On the trunk, the dermal mesenchyme of the back originates from the somites and that on the belly from the lateral plate mesoderm^2^. In the skin, the dermal mesenchymal cells acquire different fates based on their depth relative to the epithelium, with WNT signals maintaining the papillary dermis fate in the upper dermis and suppressing the acquisition of adipocyte identity, resulting in dermal adipocytes forming only in deeper dermis^3,4^. In adult skin, dermal fibroblasts decrease in number with age and are non-motile, while embryonic populations undergo extensive movement^5,6^.

Cells in the upper dermis form condensates at sites of emerging skin appendages, such as hair and feather follicles, morphologically similar to the condensates that form the mesenchymal component of intestinal villi, tooth buds and cartilaginous template of the skeleton^7–9^. These dermal condensates are formed through recruitment of mesenchymal cells by focally produced epithelial signals, notably FGF20 and SHH^6,10–12^, though autonomous coalescence of mesenchymal cells can be triggered under certain conditions without epithelial signals^6,12^. Dermal condensates have high developmental potential, and induce fate commitment and appendage growth in their overlying epithelium^13–16^. In embryonic skin, proliferation of mesenchymal cells occurs widely outside condensates^11,17^ but rarely once cells enter them^18,19^. A select, highly proliferative fibroblast population in the peri-condensate zone has been hypothesised to provide cells for condensate construction, based on cell labelling with thymidine analogues and gene expression analyses^18,20^.

Here, we delineate dermal mesenchymal cell behaviour by direct observation in developing skin, finding a signature mechanosensitive migratory mechanism that both disperses mesenchymal cells in the general dermis but also aids their recruitment into developing condensates.

## Results

### Rapid displacement of newly born mesenchymal cells in embryonic skin

To define individual cell behaviors during mesenchymal development, we analysed time-lapse confocal imaging of ex *vivo* cultured TCFLef::H2B-GFP mouse skin containing H2B-GFP labelled mesenchymal cell nuclei^21^, allowing us to track their movement throughout the cell cycle. Our observations first indicated that movement and mitosis of dermal cells occur in a planar manner in parallel to the epithelium at all depths of mesenchyme (Figure 1A and Video S1).

**Figure 1.**
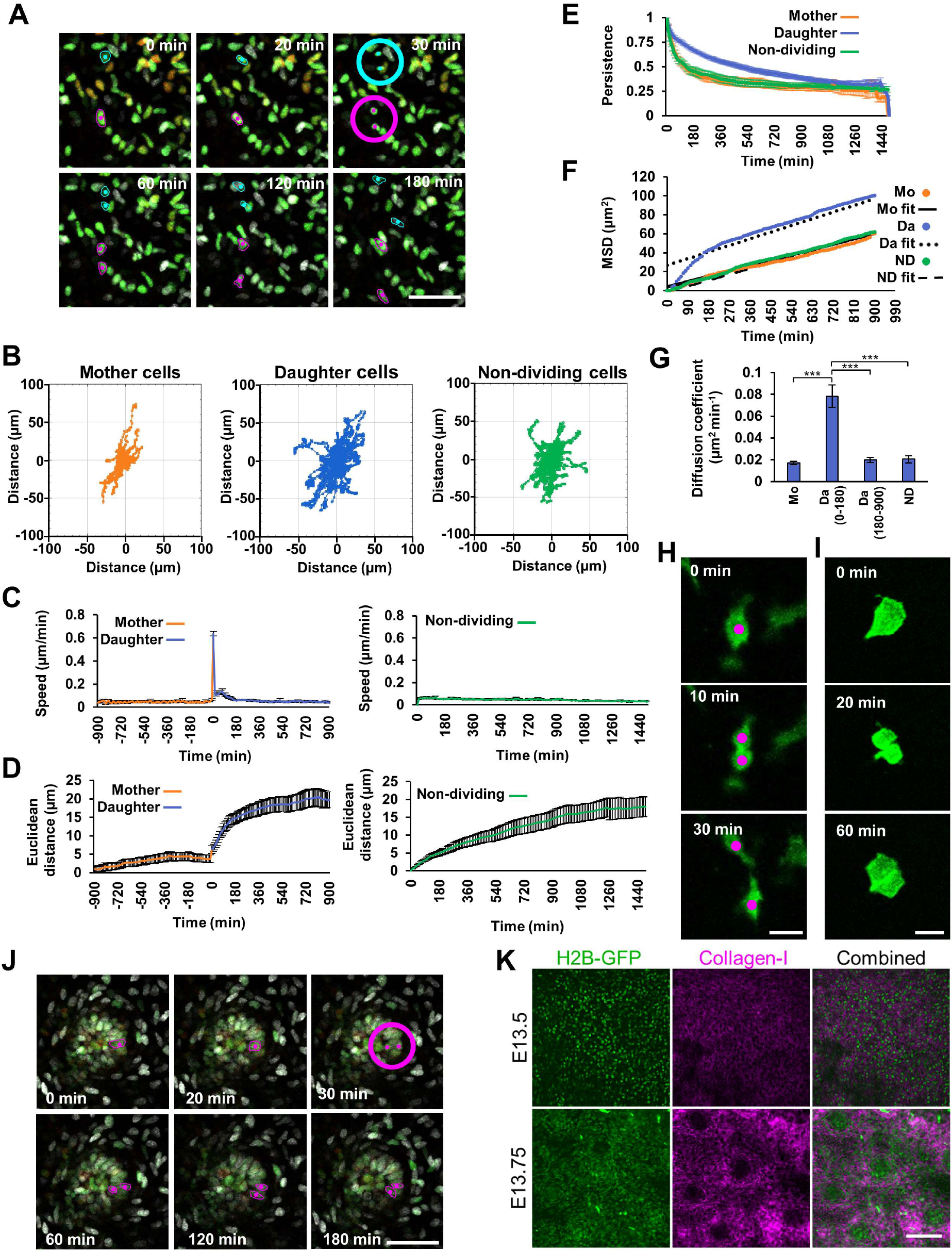
Mesenchymal mitosis is followed by rapid daughter cell displacement. **(A)** Time-lapse series of embryonic day **(E)** 13.5 TCF/Lef::H2B-GFP mouse skin explant culture. Magenta and cyan dots highlight nuclei undergoing mitosis, circles indicate the point at which cells divide, z-planes are colour-coded (red = closest to epithelium, green = middle, grey = deepest). **(B) D**iffusion plots showing the migration direction and dispersion of tracked mother (left; n=45), daughter (middle; n=89) and non-dividing (right; n=51) cells from a single skin explant. **(C-D)** Plots showing speed **(C)** and Euclidean distance travelled **(D)** by tracked dividing (left panels; point of mitosis set at time = 0) and non-dividing (right panels) cells. **(E)** Persistence (Euclidean/accumulated distance) of mother, daughter and non-dividing cells against time. **(F)** Mean squared displacement (MSD) of mother (Mo; n=45), daughter (Da; n=89) and non-dividing (ND; n=51) cells over time from a single skin explant. **(G)** Mean diffusion coefficient (the slope of the line in F) of mother cells (Mo), daughter cells (Da) up to 180 minutes after division, daughter cells after 180 minutes of division, and non-dividing cells (ND). A one-way ANOVA (p=1.03×10^-8^) and post-hoc Tukey test revealed a significant difference between newly born daughters (0-180 min) and all other groups (***p<0.001). **(H-I)** Single planes from confocal time-lapse series of a cultured E13.5 mTmG mouse skin showing (**H**) a dividing mesenchymal cell and resulting daughter cells and **(I)** a dividing peridermal cell. **(J)** Time series of mitosis in a dermal condensate. Magenta dots highlight mother and daughter nuclei. z-planes colour-coded as in **A**. **(K)** Single planes from confocal imaging of Collagen-I in E13.5 and E13.75 TCF/Lef::H2B-GFP explants. **C**, **D**, **E** and **G** time-lapse videos n=8 from 4 independent samples, mean number of dividing cells tracked per video = 42, nondividing cells tracked per video = 50. Error bars represent the standard error of the mean (SEM). Scale bar in **A** = 50 μm; scale bar in **H**-**I** = 20 μm; scale bar in **J** = 50 μm; scale bar in **K** = 100 μm

We assessed the behaviour of dividing cells and of cells that were not observed to divide within the duration of observation (hereafter referred to as non-dividing cells) in our time-lapse movies. Cells that did not divide over the 25 hour duration of the time-lapse migrated without preferred direction (Figure 1B). Occasional cells (frequency of <1/1000) migrated at high speed throughout the imaging period (Video S2), but most moved at a mean speed of 0.02 μm min^-1^ (Figure 1C). Upon and following mitosis, we observed a stereotyped daughter cell displacement, characterised by rapid separation at cytokinesis, often followed by a short pause and then a 180 minute period of rapid, persistent migration in early interphase, before returning to the behavioural characteristics of non-dividing cells (Figures 1C-1G, S1 and Video S3). As a population, the diffusion coefficient of daughter cells in this window was significantly greater, resulting in a mean Euclidean displacement ~50% greater than in the equivalent nondividing cells over the same time period (Figure 1D and 1G). To determine if this newly born cell behaviour was conserved in vertebrates, we analysed dividing mesenchymal cell behaviour in chicken skin using recombinant TAT-Cre induced Chameleon chicken cells^11^ (Figure S2A and Video S4) in explant skin culture. We observed a similar increase in speed and displacement of newly born cells, though we did not detect an increased persistence of movement (Figures S2B-S2G). In line with these short-term observations, longer term lineage tracing of cells derived from the somites or the surface ectoderm using TAT-Cre induction in the Chameleon chicken line over four days revealed the formation of elongated contiguous keratinocyte clones in the epidermis running dorsolaterally, while mesenchymal clones, although elongated and running dorsolaterally, were not contiguous (Figure S2H), consistent with dispersal of mesenchymal cells immediately upon mitosis. Similarly, when we transplanted a somite from a tdTomato (TPZ) chicken into a GFP chicken, we observed a dorsolateral streak of tissue expansion originating from the somite (Figure S2I), reflecting a favoured net direction of cell displacement in the growing embryo.

To assess individual cell shape through mitosis we induced sparse labelling of membrane localised MARCKS-EGFP in mesenchymal, periderm and basal epidermal cells using the *R26R-mTmG* mouse line^22^. We induced labelling using cell permeable TAT-Cre in E13.5 skin, then cultured and confocal imaged for 16 hours. We noted that dermal mesenchymal cells typically have planar cell projections, running parallel to the epithelium. These projections are often, but not always, lost upon cell division, with rounding of the cells typically occurring as they enter mitosis (Figure 1H). However, several instances of retention of major cellular processes, followed by migration along their tracks, were evident (Video S5). The behaviour we observed in mesenchymal cells was distinct from that in epithelia, where we observed divisions of periderm and of basal epidermal cells which produced adjacent daughter cells (Figure 1I and Video S6), in agreement with the contiguous epithelial clones we observed in chicken embryos over a longer timescale (Figure S2G).

Mesenchymal fibroblasts contribute to hair follicles by forming dermal condensates, precursors to the dermal papilla and the follicle’s peripheral dermal sheath^23^. Though the embryonic dermal mesenchyme is highly proliferative, cells in the dermal condensates are long recognised as being largely quiescent^10,18,20^. However, in our imaging experiments we observed some mitoses taking place within dermal condensates (Figure 1J and Video S7). In sharp distinction to the general dermis, these mitoses resulted in very limited cell displacement, producing a pair of daughter cells that remained in close proximity to one another, suggesting that the local extracellular environment influences the behaviour of newly born cells in the mesenchyme. To assess this, we immunostained embryonic mouse skin, and found that Collagen-I was absent from the dermal condensates, revealing their distinct extracellular matrix environment (Figure 1K). We also observed that Collagen-I is increased around the periphery of the dermal condensates in a ring, possibly displaced by cell compaction into the dermal condensate.

### Rapid movement of the newly born mesenchymal cells increases dermal condensate entry

Dermal mesenchymal cells are drawn to condensates by soluble signals, notably FGF20 and SHH, from the overlying epithelial placodes^6,10^. Consistent with an epithelial recruitment influence, we observed that dermal condensates form from the epithelium downwards (Figure S3). In agreement with previous findings^18,20^, our imaging experiments found that cells within the first 180 minutes post-division were more likely to enter a dermal condensate than nondividing cells, with the rate of entry declining to that of the general interphase population thereafter (Figure 2A and Table S1).

**Figure 2.**
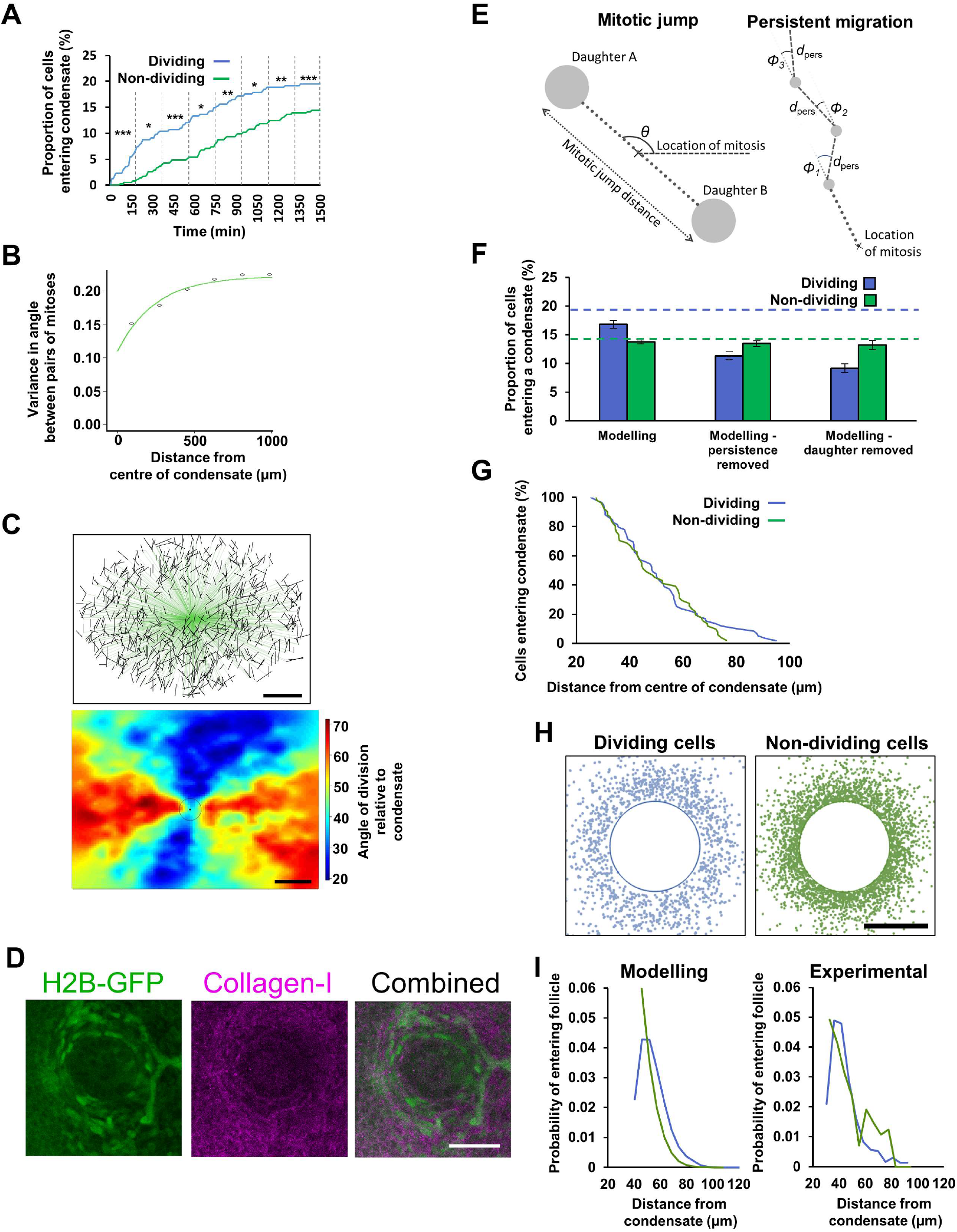
Newly born mesenchymal cells have increased probability of dermal condensate entry. **(A)** Timing and proportion of tracked dividing and non-dividing cells entering the condensates as a cumulative percentage. Vertical dashed lines and asterisks indicate time windows and significance levels for a Fisher’s exact test (Table S1). Time 0 is the point of mitosis for dividing cells. Cells from 0 to 180 minutes post-mitosis have an increased rate of condensate entry compared to all other cells (*p < 0.05, **p < 0.01, ***p<0.001). **(B)** Spatially-dependent variance (variogram) in mitosis angles. Lower variance is found closer to the follicle (distance = 0), 51% of the variance in the mitoses direction is explained by the distance from the follicle (Monte Carlo probability p <0.001). **(C)** Upper: Cell division angles relative to their nearest condensate for a single time-lapse sequence. Black lines indicate direction of mitoses, green lines connect division events to the condensate centre, from which angles relative to the condensate were calculated. Lower: heat map of the cell division angles relative to nearest dermal condensate for the same dataset (black circle = centre). **(D)** Single plane confocal images of Collagen-I stained E15.5 TCF/Lef::H2B-GFP skin. Peri-condensate nuclei align with long axes tangential to the condensate. **(E)** Schematic of division producing daughters undergoing post-mitotic displacement (left panel). Daughters A and B are equidistant from the location of mitosis. Daughter A moves away from the location of mitosis in the diametrically opposite direction (θ) to B. Possible trajectory of daughter A after division (right panel). The dotted line shows the direction of mitotic jump, followed by a persistent random walk (dashed lines), from which angles of migration relative to the direction of the mitotic jump are represented by Φi, and the distance travelled in each direction is represented by d_pers_. **(F)** Plot showing the percentage (+/- SEM; n=8) of simulated dividing and non-dividing cells entering a condensate using modelling based on experimentally-determined parameters. Dashed blue and green lines indicate dividing and non-dividing entering proportions from experimental data. **(G)** Plot showing percentage of dividing and non-dividing cells entering a condensate against their initial position relative to the condensate centre from experimental data. Lineages with recent divisions can be recruited from further away. **(H)** Initial locations of dividing (left) and non-dividing (right) agents that ultimately enter a condensate, from simulation. **(I)** For cells entering follicles, the probability density of entry from a given starting distance for dividing (blue) and non-dividing (green) cells - modelled (left) and experimental (right). Scale bars in **C** = 100 μm; scale bar in **D** = 50 μm; scale bar in **H** = 50 μm.

We next examined whether the dermal condensate influences the orientation of nearby cell divisions. We observed that the mitosis direction was affected by the presence of a condensate up to a mean of 295 μm from its centre (Figure 2B). Performing a permutation analysis of mitosis angles we found that, at distances closer than 300 μm to the follicle, the distribution of mitosis angles cannot be explained by a random process, indicating that condensates influence division angle within this radius. We also observed that mitoses occur at a tangent to the condensate perimeter (Figure 2C and Video S8). By immunostaining E15.5 mouse skin, in which condensates are well-formed, we found that cells were elongated and aligned within the collagen ring surrounding the condensate, suggesting that the extracellular environment surrounding the condensate influences the orientation of cells and the direction of mitosis (Figure 2D). Therefore, dermal condensates influence the division angle of mesenchymal cells in their immediate proximity. However, dividing cells do not orientate directly towards the condensate, and thus this effect does not explain the higher proportion of dividing cells ultimately entering the condensate (19.4%), compared to non-dividing cells (14.3%; Figure 2A).

In order to investigate the mechanisms underlying the observed recruitment bias in favour of newly born mesenchymal cells to condensates, we built an agent-based mathematical model (Figure 2E and appendix). Dermal fibroblast cells (the agents) were initialised uniformly at random on a square domain (representing the dermis) with periodic boundary conditions that contained preformed condensate recruitment sites with fully absorbing boundary conditions.

Cells encountering the condensate are immediately absorbed and removed from the simulation, consistent with our findings that cells are not observed exiting established condensates. Cell movement on the domain is an off-lattice diffusive random walk, with migration parameters being those directly measured from the non-dividing mesenchymal cell population (Figure 1F). To model mitosis, cells were selected at random, then simulated to undergo a proliferation event forming two daughters, which then underwent mitotic displacement over 180 minutes according to our measured cell movement parameters (Figure 2E and see appendix). In this model, newly born mesenchymal cells have a greater likelihood of condensate entry than non-dividing cells as a result of both cell number increase (through division) and the increased displacement of the resulting daughters, making them more likely to encounter a condensate (Figure 2F). Our modelling predicted that the probability of dividing cells being recruited from short distances would be lower than for non-dividing cells but that they could be recruited from a greater distance, as suggested by our cell tracking data (Figure 2G) and confirmed by plotting the probability density (Figures 2H and 2I).

### Mitotic displacement of mesenchymal cells is locally coordinated in intact embryos

In our initial imaging of skin cultures we noted that mitotic angles tended to be aligned with one another across the field of view (Figure 3A). To determine whether this observation was unique to ex *vivo* culture, we imaged intact E13.5 TCFLef::H2B-GFP embryos using 2-photon microscopy (Figure 3B) and tracked mesenchymal cell behaviour for 2 hours. Here we observed the same stereotyped planar mitotic displacement mechanism (Figures 3C and 3D). Global mesenchymal cell division angles were not aligned with the cardinal axes of the embryos (Figure 3E). However, spatial statistical analysis confirmed that contiguous zones of coherent, aligned cell divisions were present (Figure 3F), with areas of correlated mitoses accounting for around 80% of the total area in our images. This suggests that the influence on mitosis angle that is global in our 2-dimensional skin cultures is instead more regional in intact and growing embryos, perhaps reflecting their more complex topology.

**Figure 3.**
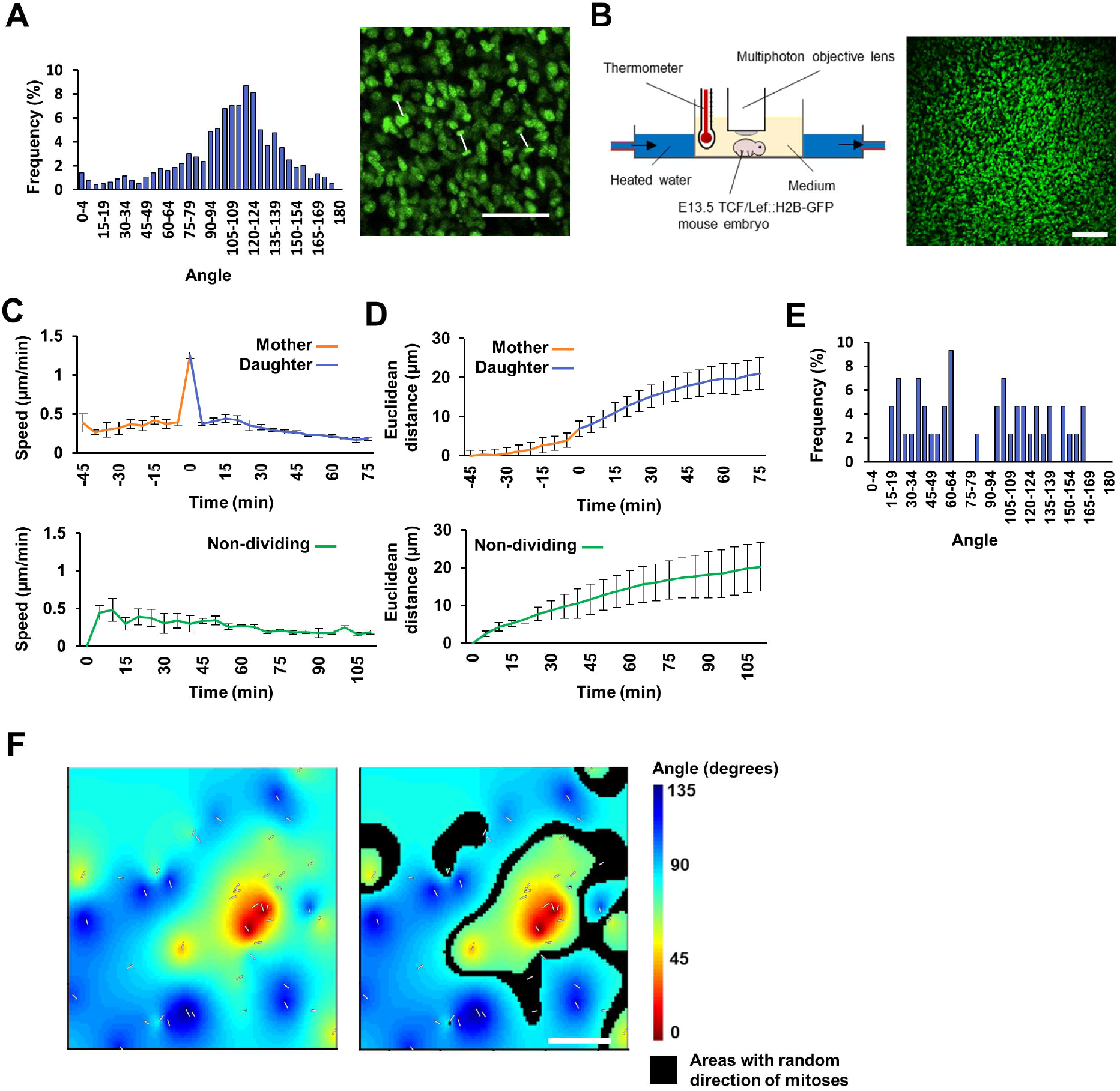
Mitotic orientations are locally aligned in embryonic mesenchyme. **(A)** Distribution of mitotic angles (n=1135) in a TCF/Lef::H2B-GFP skin explant culture (left panel) and representative frame showing coherence of mitotic orientations (right panel). Daughter nucleus pairs are connected by white lines, showing angle of mitosis. **(B)** Schematic of whole embryo culture and imaging, and corresponding image of the trunk skin of an E13.5 TCF/Lef::H2B-GFP embryo. **(C-D)** Speed **(C)** and Euclidean distance travelled **(D)** by tracked dividing and non-dividing cells (whole embryo time-lapse from 3 independent samples, average number of divisions tracked/video = 65, non-dividing cells tracked/video = 50. In dividing cell plots, time 0 = mitosis). **(E)** Distribution of angles (n=43) of division from TCF/Lef::H2B-GFP whole mouse embryo imaging. **(F)** Spatial distribution of mitotic angles in **E** (white overlaid lines), with heat map showing areas where angles are correlated. Right panel shows areas (overlain in black) containing no significant local correlation (i.e. mitotic angles are random). The randomness ratio (proportion of black area to total area) calculated at a significance level of 0.01, ranged from 16% to 24% between the fields of view analysed. Error bars represent SEM. Scale bar in **A** = 50 μm; scale bar in **B** = 100 μm; scale bar in **F** = 200 μm.

### The direction and speed of newly born mesenchymal cell migration are mechanosensitive

Having observed correlation of mitotic angles in small domains surrounding the dermal condensates and in larger zones in their absence, we hypothesized that they were likely influenced by mechanical forces. To test this, we compared embryonic mouse skin that had been relaxed fully in culture (Figure S4A) with skin stretched to approximately twice its length along a single axis (Figure S4B). Relaxed skin carried nuclei with a generally oval morphology and no alignment of orientation, while stretched showed alignment of the long axis of the mesenchymal nuclei with the applied tension (Figures 4A, S4C and S4D), though more pronounced for a lateral (dorsal-ventral) stretch than an equivalent axial (anterior-posterior) stretch. A lateral stretch also caused an increase in average nucleus length (Figure 4B). Immunofluorescent detection of Collagen-I confirmed that fibres of the extracellular matrix had aligned with the direction of stretch (Figures S4E and S4F).

**Figure 4.**
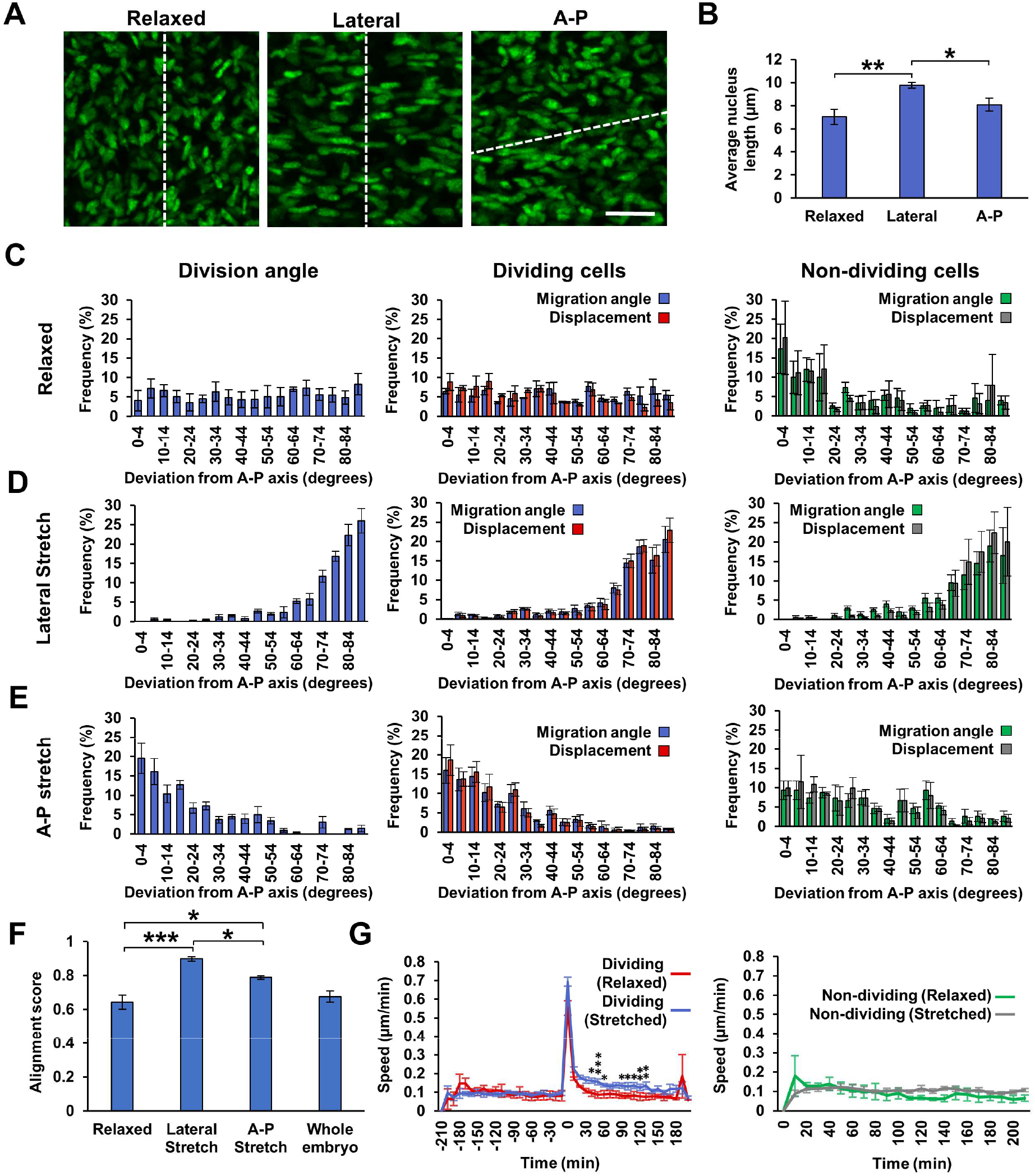
Orientation of mesenchymal mitosis and newly born cell migration by mechanical cues. **(A)** E13.5 TCF/Lef::H2B-GFP mesenchymal nuclei in skin in relaxed and stretched states. Skins were stretched to approximately twice their length along a single axis. White dashed line indicates the anterior-posterior (A-P) axis. **(B)** Nucleus length from relaxed (n=3), laterally stretched (n=4) and A-P stretched (n=3) skins. **(C-E)** Plotted deviations of cell division angles (left), and migration angle of dividing (middle) and non-dividing (right) cell migration angle from the A-P axis and total cell displacement for each angle in skins that were **(C)** relaxed (n=3, average number of dividing cells tracked/video = 67, non-dividing cells tracked/video = 50), **(D)** stretched laterally, (n=4, average number of dividing cells tracked/video = 72, non-dividing cells tracked/video = 50) and **(E)** stretched along the A-P axis (n=3, average number of dividing cells tracked/video = 79, non-dividing cells tracked/video = 50). **(F)** Global alignment scores across entire skin of mitosis angles from relaxed skins (n=3), laterally stretched skins (n=4), A-P stretched skins (n=3) and whole embryos (n=3). **(G)** Speed of tracked dividing (left panel; time 0 = point of mitosis) and non-dividing (right panel) cells in skins that were relaxed (n=3, average number of dividing cells tracked/video = 67, non-dividing cells tracked/video = 50) or stretched (n=7, average number of dividing cells tracked/video = 78, non-dividing cells tracked/video = 50; *p < 0.05, **p < 0.01, ***p<0.001). Error bars represent SEM. Scale bar in **A** = 20 μm.

We tracked cell division and migration in relaxed and stretched E13.5 skin cultures. In relaxed skin, cell migration and mitotic displacement occurred with the same characteristics as observed earlier (Figures 1C, 1D and S5), but with no preferred angle of mitotic division or migration in dividing cells. Non-dividing cells exhibited a slight preference for anterior-posterior movement (Figure 4C). In stretched skin, mitotic displacements and the migration of the newly born and the non-dividing cells aligned with the direction of applied stretch (Figures 4D, 4E, and Video S9). Mitotic displacements in chicken skins also aligned with the direction of tension (Figure S4G). As observed for nuclear elongation, a lateral stretch was more effective in aligning mitotic angles than an equivalent axial stretch (Figure 4F), and relaxed skins showed a similar global alignment score to that from our whole embryo imaging analysis (Figures 3E and 4F). Strikingly, we found a unique responsiveness of newly born cells to tissue strain, in which daughter cells move more rapidly in stretched skin compared to relaxed skin, while mother and non-dividing cells did not change their speed of movement. This reveals that newly born mesenchymal cells enter a more mechanosensitive state in early G1 phase, responding to tissue strain with increased migration rates (Figure 4G).

## Discussion

We report, from direct observation, a stereotypical migratory behaviour of early G1 phase mesenchymal cells that drives their displacement in tissue growth and patterning. These cells separate rapidly at mitosis and over the next 180 minutes migrate with greater speed and persistence than their peers further into G1 and S phases. This rapid migratory behaviour is not a continuation of cytokinesis, but instead the cells move rapidly and engage in one or more saltatory leaps in the 180 minutes after mitosis. Cell division and movement is planar, occurring almost entirely in parallel with the epithelium. This stability of mesenchymal cell vertical positions relative to the epithelium allows positional signals, such as epithelial WNT, to direct the appropriate differentiation of distinct layers of the dermis^1,24^.

The behaviour of mesenchymal cells at mitosis is highly mechanosensitive, orienting division and migration along lines of tension in the tissue. This matches the long-recognised orientation of dividing cells in epithelial sheets, which also align their plane of division perpendicular to orientation of tissue tension^25^. Epithelial cells remain adjacent or close to one another after division, leading to the formation of contiguous epithelial clones indicative of the mode of tissue growth, such as the sharply demarcated lines of Blaschko in the dorsal skin of mammals^19,26^. However, in the mesenchyme cell division is followed by directed dispersal of newly born daughters, immediately spreading them across the tissue. This mechanism will effectively fill gaps^27^, for example through non-uniform skin growth or stretch, to maintain a uniform cell density. In addition, this rapid displacement of sister cells enforces the disruption of mesenchymal cell clones, preventing formation of mesenchymal condensates through clonal expansion. This permits the construction of condensates through recruitment using soluble developmental signals instead^6,10,12^.

Previous work identified an increased frequency of daughter cells from recent cell divisions within mesenchymal condensates^18,20^. This result is explained by our observations as resulting from the increased dispersal rate of newly born cells compared to their mid and late G1, and S phase peers, making them more likely to encounter recruitment signals. Our simulations show that this observed behaviour alone, without a need to invoke spatially heterogeneous proliferation nor a cell cycle phase specific sensitivity to placode-derived recruitment signals, is sufficient to account for the increased representation of products of recent cell divisions in dermal condensates.

The cell behaviours we observe in general dermis are highly distinct from those in the mesenchymal condensate, where the extracellular matrix is depleted. Future work will address the mechanistic basis of this phenomenon, its generality in the body’s diverse mesenchymal populations, the relationship between intracellular events and extracellular matrix interaction and sensing, and whether orientation of mesenchymal cell division and interphase migration can be modulated by extracellular signals to achieve polarised tissue growth.

---

**Figure S1.**
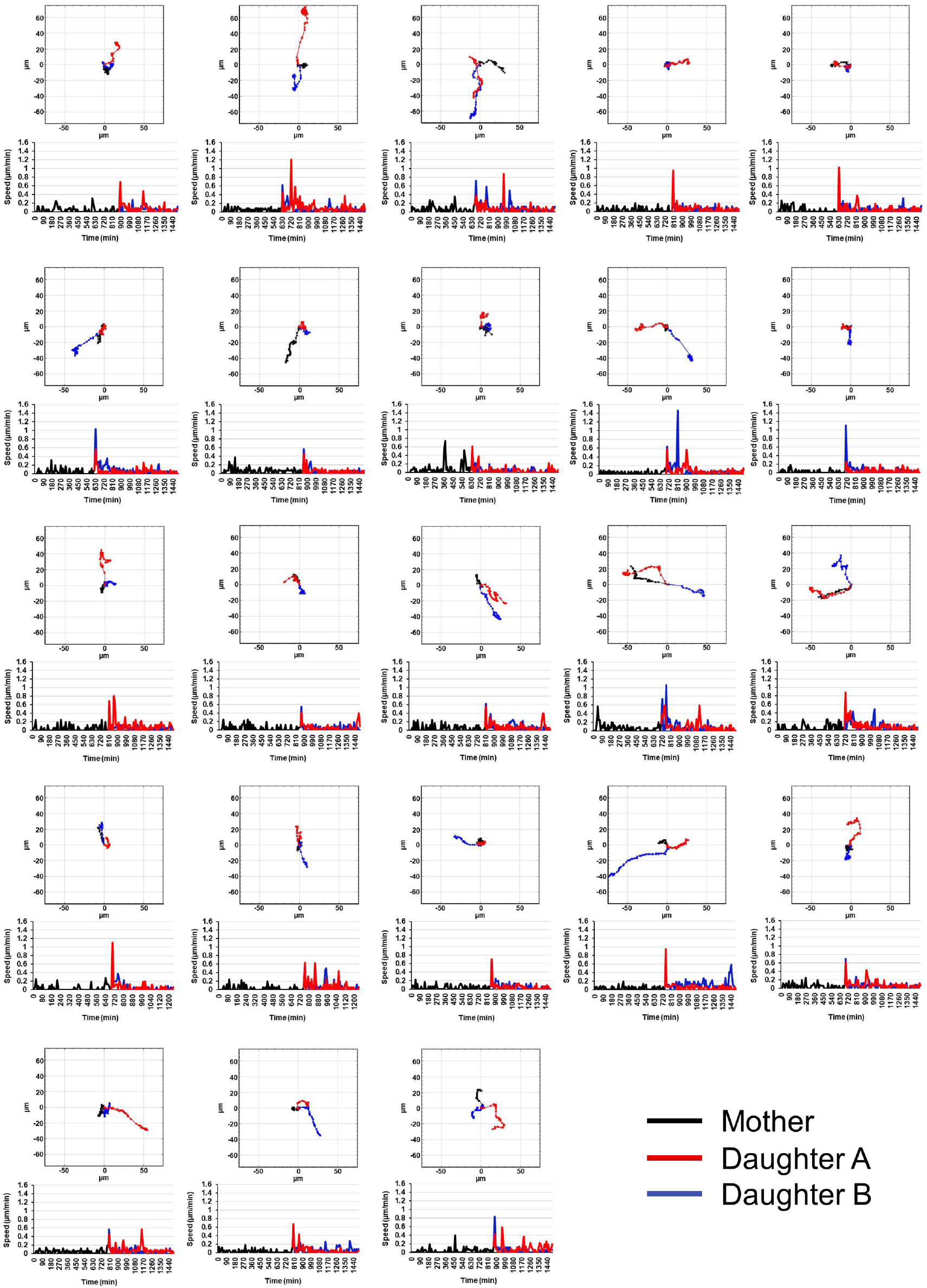
Newly born daughter cells migrate extensively after division. Mitosis plots mapping mother and daughter tracks before and after cell division, with corresponding cell speed plots over the timecourse of imaging below. Tracks were chosen from cells in which cell division took place within a timepoint of 40-60% through the imaging, and both daughter cells were tracked until the end of the video.

**Figure S2.**
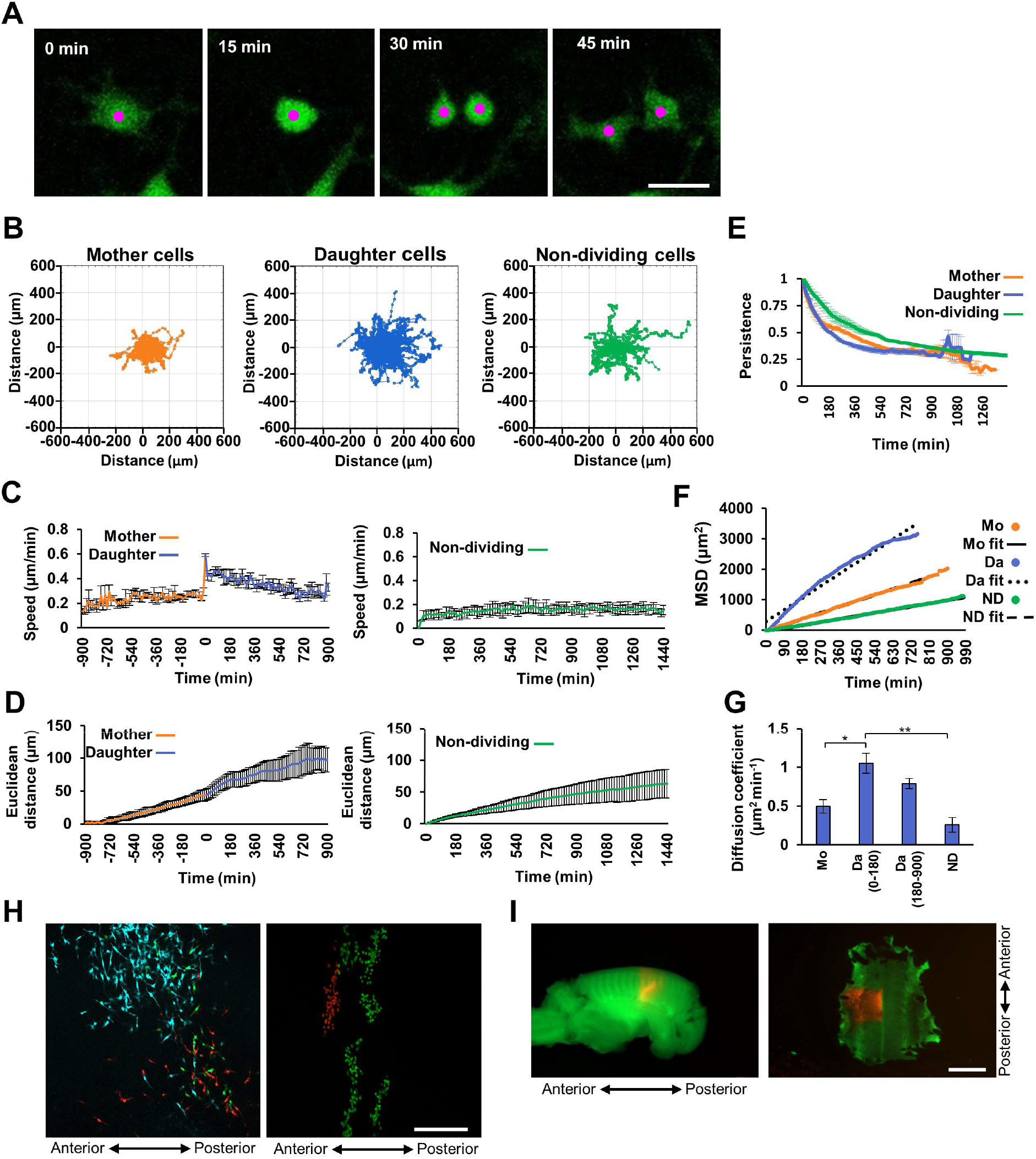
Newly born mesenchymal cells exhibit rapid migration in embryonic chicken skin. **(A)** Single z-planes from confocal time-lapse series of an E6 Chameleon chicken skin explant culture treated with cell permeant TAT-Cre and cultured for 24 hours. Magenta dots highlight mother and daughter cells. **(B)** Diffusion plots showing the migration direction and dispersion of tracked mother (left; n=62), daughter (middle; n=124) and non-dividing (right; n=50) cells from a single skin explant. (C-D) Speed **(C)** and Euclidean distance travelled **(D)** of dividing (left panels; time 0 = point of mitosis) and non-dividing (right panels) cells from Chameleon chicken skin explants. **(E)** Persistence (Euclidean/accumulated distance) of mother, daughter and non-dividing cells against time. **(F)** Plot showing mean squared displacement (MSD) of mother (Mo; n=62), daughter (Da; n=124) and non-dividing (ND; n=50) cells over time from a single skin explant. **(G)** Mean diffusion coefficient (slope of line in F) of mother cells (Mo), daughter cells (Da) up to 180 minutes after division, daughter cells after 180 minutes of division, and non-dividing cells (ND). A one-way ANOVA (p=0.002) and post-hoc Tukey test revealed a significant difference between daughters (0-180 min) and mother cells, and daughters (0-180 min) and non-dividing cells (*p<0.05, **p<0.01). **(H)** Confocal images of clonally labelled dermal and epithelial cells in E7 Chameleon skin explant culture following somite and overlying epithelium injection of cell permeant TAT-Cre at E3. Left panel, mesenchymal clones; right panel epithelial clones. **(I)** Tissue derivatives of a tdTomato somite (TPZ transgenic line) transplanted into a CAG-GFP host embryo, at E6.5. Left panel, intact embryo; right panel, isolated skin. **C**, **D**, **E** and **G** time-lapse videos n = 3 from 3 independent samples, mean number of dividing cells tracked per video = 60, mean number of non-dividing cells tracked per video = 91. Error bars represent SEM. Scale bar in **A** = 20 μm; scale bar in **H** = 200 μm. scale bar in **I** = 2 mm.

**Figure S3.**
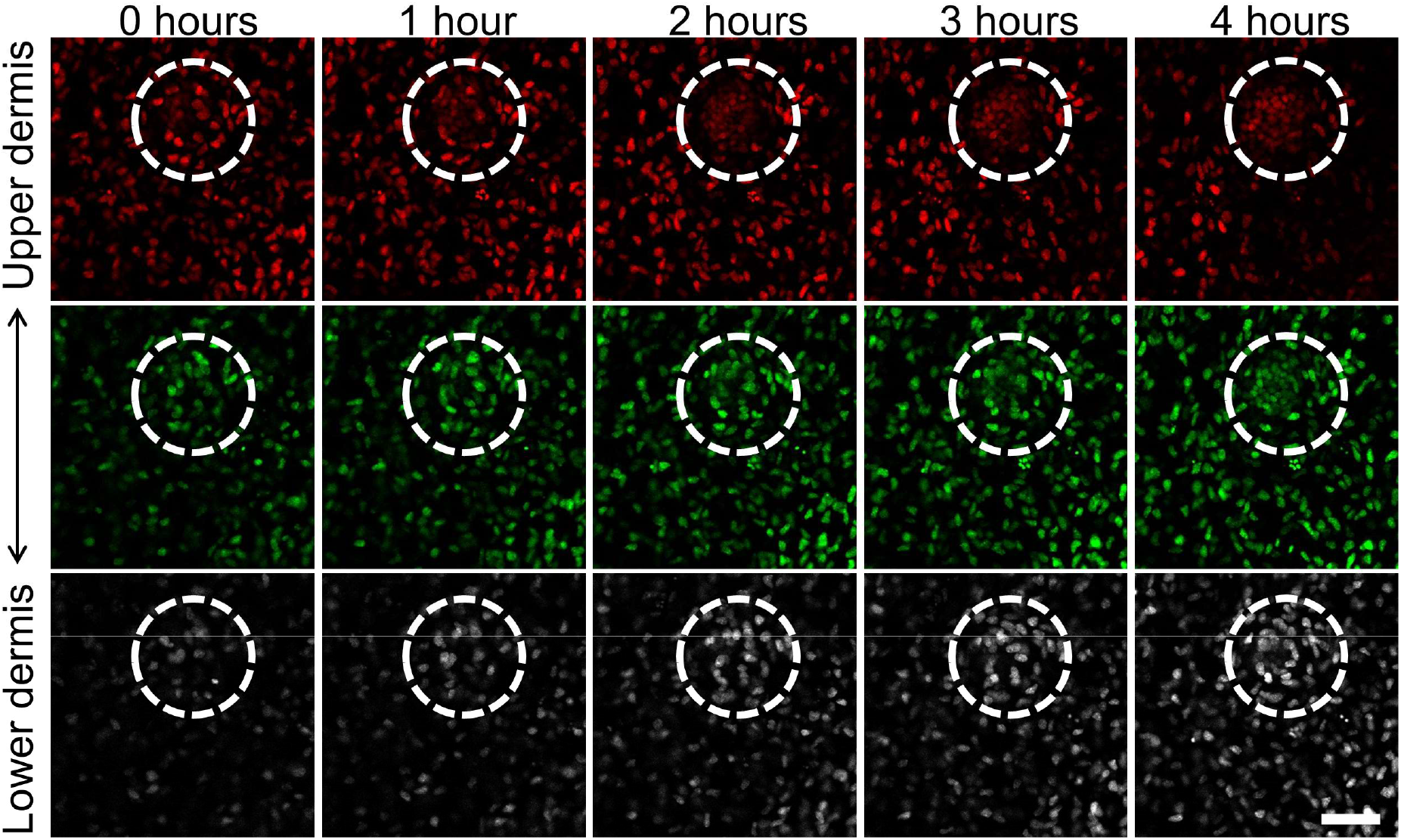
Dermal condensates form from the epithelium downwards. Depth-coded single planes from time-lapse confocal imaging of E13.5 TCF/Lef::H2B-GFP skin explants during dermal condensate formation. Red = plane closest to epithelium, green = middle plane, grey = plane furthest from epithelium. White dashed line outlines condensate location. Scale bar = 50 μm.

**Table S1.** Statistical comparisons of rate of cell entry to dermal condensates in 180 minute windows, for dividing and non-dividing cells. Newly born daughter cells have a higher rate of condensate entry compared to non-dividing cells in their first 180 minutes.

| Time Window | Dividing cells entering condensate | Dividing cells not entering condensate | Non-dividing cells entering condensate | Non-dividing cells not entering condensate | Fisher's exact test p-values |  |
| --- | --- | --- | --- | --- | --- | --- |
|  |  |  |  |  | Dividing frequency (0-180) vs non-dividing frequencies | Dividing frequency (0-180) vs dividing frequencies |
| **0-180** | 21 | 287 | 3 | 350 | 0.0000328* |  |
| **190-370** | 11 | 276 | 11 | 339 | 0.0305094* | 0.144831 |
| **380-560** | 5 | 271 | 5 | 334 | 0.0005216* | 0.0041089* |
| **570-750** | 9 | 262 | 9 | 325 | 0.0148993* | 0.0624126 |
| **760-940** | 7 | 255 | 7 | 318 | 0.0058911* | 0.0308804* |
| **950-1130** | 5 | 250 | 8 | 310 | 0.0126678* | 0.0076809* |
| **1140-1320** | 1 | 249 | 6 | 304 | 0.0029695* | 0.0000354* |
| **1330-1500** | 1 | 248 | 2 | 302 | 0.0000497* | 0.0000363* |

\*denotes $p < 0.05$ threshold

**Figure S4.**
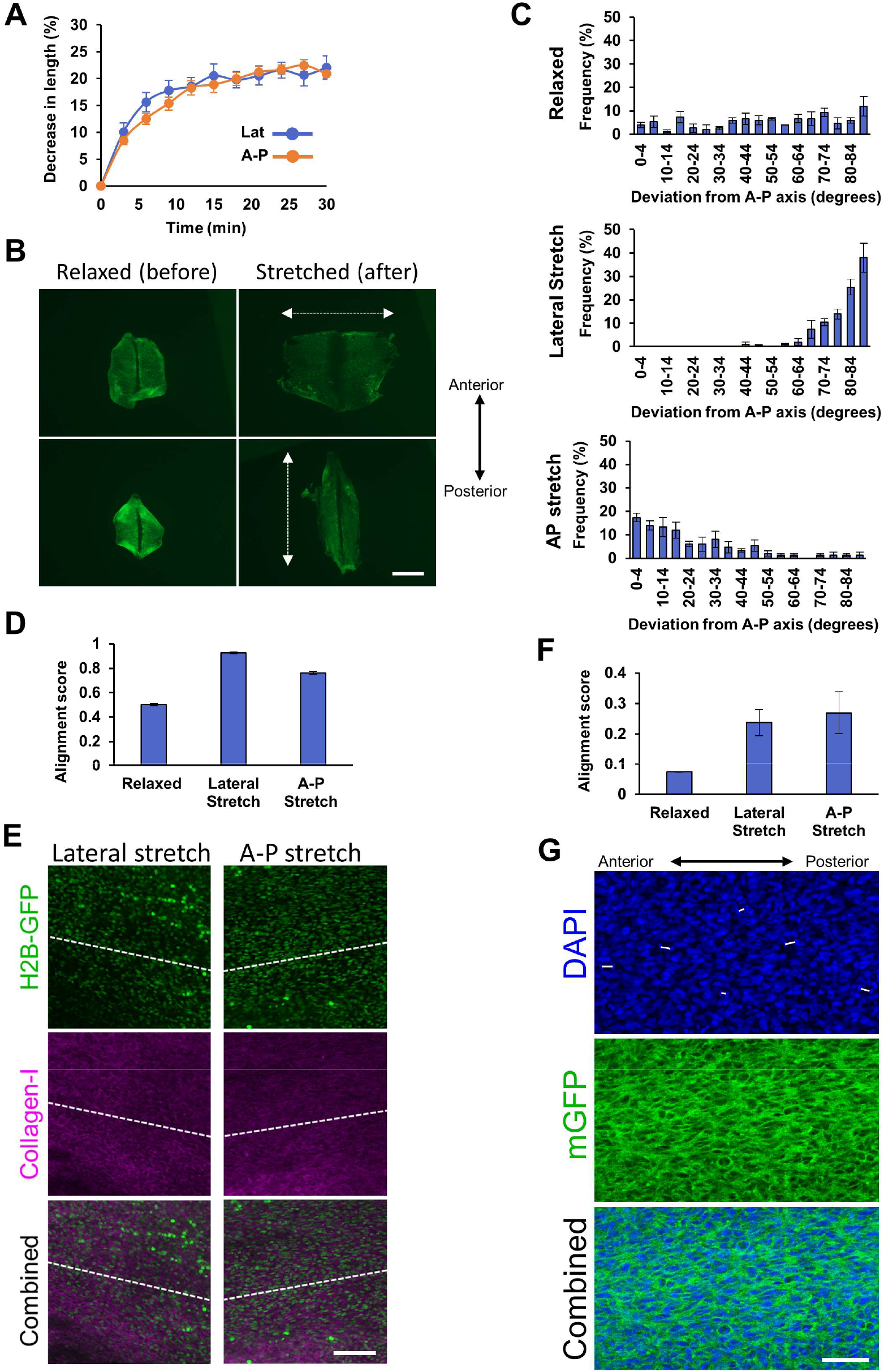
Nuclei and Collagen-I align with mechanical strain. **(A)** Plot showing the percentage decrease in anterior-posterior (A-P) and dorsal-ventral (Lat) length of mouse skin explants in a 30 minute period (n=5) when suspended freely in culture medium. **(B)** Images of E13.5 TCF/Lef::H2B-GFP mouse skin explants before and after a lateral stretch (upper panels) or a stretch along the anterior-posterior (A-P) axis of the embryo (lower panels). White dashed arrows show direction of stretch. **(C)** Nucleus orientation angle for relaxed (n=3; upper), lateral stretched (n=4; middle) and A-P stretched (n=3; lower) skins. **(D)** Alignment score of nucleus orientation angle from relaxed (n=3), laterally stretched (n=4) and A-P stretched (n=3) skins. **(E)** Single planes from confocal imaging of Collagen-I immunofluorescence in E13.5 TCF/Lef::H2B-GFP skin explants in stretched states. Dashed white line indicates direction of stretch. **(F)** Alignment scores of collagen fibres from relaxed, laterally stretched and A-P stretched skins. **(G)** Single planes from confocal imaging of an A-P stretched E7 membrane GFP (mGFP) chicken skin. Daughter nucleus pairs are connected by white lines, showing coherent angles of mitosis aligned with applied tension. Error bars represent SEM. Scale bar in **B** = 2 mm; scale bar in **E** = 100 μm; scale bar in **G** = 50 μm.

**Figure S5.**
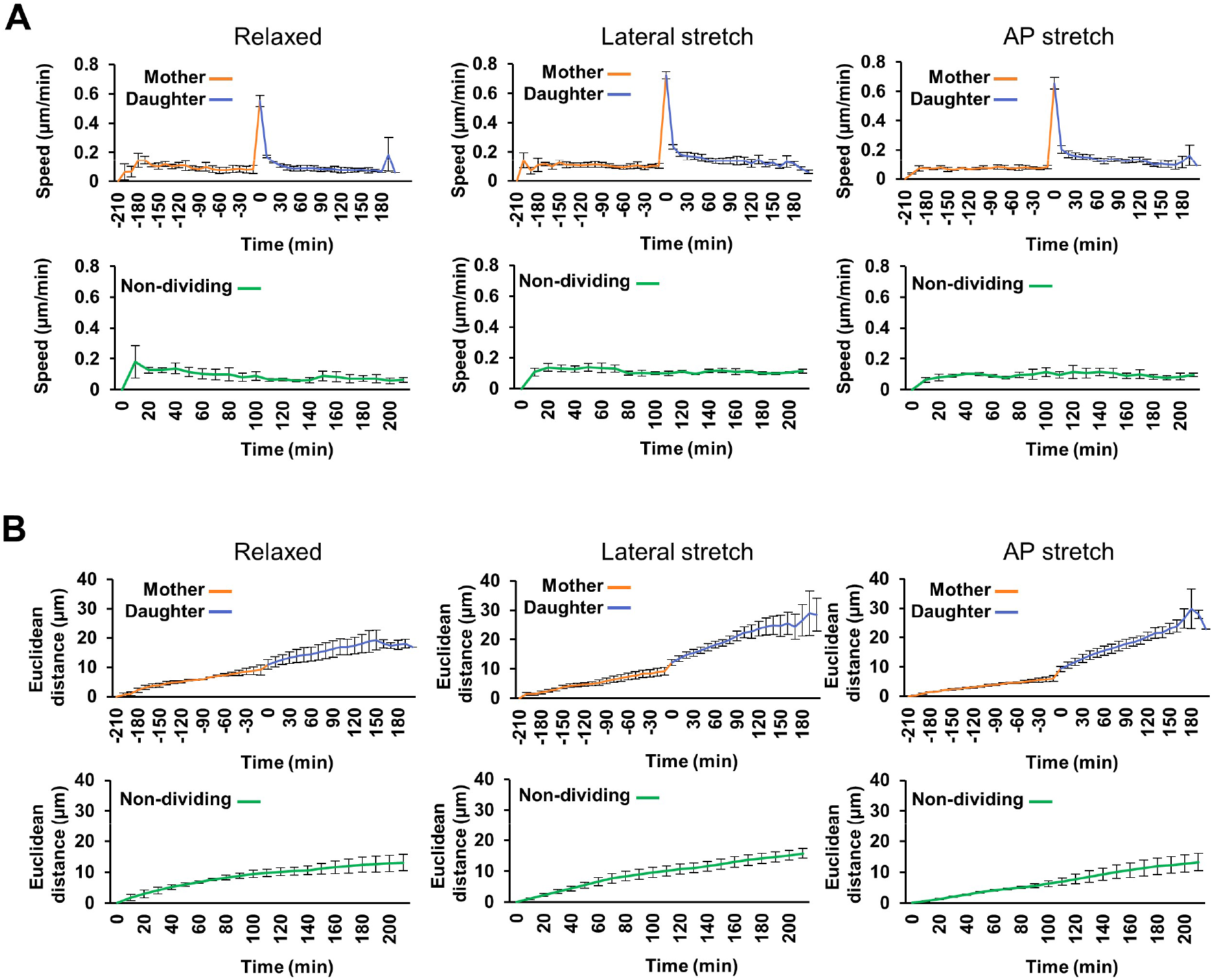
Newly born cells in stretched skin show increased speed and displacement. **(A-B)** Speed **(A)** and Euclidean distance travelled **(B)** of tracked dividing (top panels; time 0 = point of mitosis) and non-dividing (bottom panels) cells in skins that were relaxed (n=3, average number of dividing cells tracked/video = 67, non-dividing cells tracked/video = 50), stretched laterally, (n=4, average number of dividing cells tracked/video = 72, non-dividing cells tracked/video = 50) and stretched along the A-P axis (n=3, average number of dividing cells tracked/video = 79, non-dividing cells tracked/video = 50). Error bars represent SEM.

**Video S1 – Time-lapse imaging of dividing cells in TCF/Lef::H2B-GFP mouse skin**

Confocal time-lapse imaging of a cultured E13.5 TCF/Lef::H2B-GFP skin explant. Nuclei in different z-planes are colour-coded (red = closest to epithelium, green = middle, grey = plane furthest from epithelium). Magenta, cyan and red dots highlight nuclei undergoing mitosis; magenta, cyan and red circles indicate the point at which cells divide.

**Video S2 – Fast moving cell in dermis**

Confocal time-lapse imaging of a cultured E13.5 TCF/Lef::H2B-GFP skin explant. Nuclei in different z-planes are colour-coded (red = closest to epithelium, green = middle, grey = plane furthest from epithelium). A fast moving cell is highlighted by a magenta dot and circle.

**Video S3 – Daughter cells pause after cytokinesis before making large migration movements**

Confocal time-lapse imaging of a cultured E13.5 TCF/Lef::H2B-GFP skin explant. Nuclei in different z-planes are colour-coded (red = closest to epithelium, green = middle, grey = plane furthest from epithelium). Magenta and cyan dots highlight dividing cells that pause after division before moving rapidly. Magenta and cyan circles indicate the point at which cells divide.

**Video S4 – Time-lapse imaging of dividing cells in chicken skin**

Confocal time-lapse imaging of sparsely labelled and cultured E6.5 Chameleon chicken skin explant. Magenta, cyan and red dots highlight cells undergoing mitosis, magenta, cyan and red circles indicate the point at which cells divide. Note that the entire cell volume is labelled with these cytoplasmic fluorescent proteins.

**Video S5 – Time-lapse imaging of dividing mesenchymal cells in mTmG mouse skin**

Confocal time-lapse imaging of cultured E13.5 mTmG skin explant. Magenta and cyan dots highlight mesenchymal cells undergoing mitosis, magenta and cyan circles indicate the point at which cells divide. White arrowhead points to a cell process that persists through mitosis.

**Video S6 – Time-lapse imaging of dividing epithelial cell in mTmG mouse skin**

Confocal time-lapse imaging of cultured E13.5 mTmG skin explant, showing an example of a peridermal cell undergoing mitosis. The daughter cells remain in contact with one another.

**Video S7 – Time-lapse imaging of cells dividing within a dermal condensate in TCF/Lef::H2B-GFP mouse skin**

Confocal time-lapse imaging of a cultured E13.5 TCF/Lef::H2B-GFP skin explant. Nuclei in different z-planes are colour-coded (red = closest to epithelium, green = middle, grey = plane furthest from epithelium). Magenta and cyan dots highlight nuclei undergoing mitosis within a condensate, magenta and red circles indicate the point at which cells divide. In contrast to the general dermal mesenchyme, dividing cells within dermal condensates produce daughters that remain as neighbours.

**Video S8 – Time-lapse imaging of peripheral cells dividing tangentially around a dermal condensate in TCF/Lef::H2B-GFP mouse skin**

Confocal time-lapse imaging of a cultured E13.5 TCF/Lef::H2B-GFP skin explant. Nuclei in different z-planes are colour-coded (red = closest to epithelium, green = middle, grey = plane furthest from epithelium). Magenta and cyan dots highlight nuclei undergoing mitosis around a condensate, magenta and red circles indicate the point at which cells divide. Cells that are peripheral to the condensate have elongated nuclei running at a tangent to the condensate, and divide along this axis.

**Video S9 – Time lapse imaging of cell divisions aligning with stretch in TCF/Lef::H2B-GFP mouse skin**

Confocal time-lapse imaging of a cultured E13.5 TCF/Lef::H2B-GFP skin explant stretched along the dorsolateral axis. Magenta circles highlight nuclei undergoing mitosis, in which the division angle aligns with the direction of applied stretch.

## Materials and methods

### Animal lines

TCF/Lef::H2B-GFP embryos were obtained by crossing male hemizygous TCF/Lef::H2B-GFP mice on a FVB/N genetic background with female FVB/N mice. mTmG mice were on a C57Bl/6JCrl background.

Hyline non-transgenic, transgenic green fluorescent (CAG-GFP), TPZ, Chameleon and membrane GFP reporter chicken eggs were obtained from the Roslin Institute National Avian Research Facility (NARF). Chicken eggs were incubated vertically at 37.8°C.

### Tissue explant culture and stretching experiments

Dorsolateral skin was dissected from embryos in PBS and attached onto MF-Millipore™ membrane filters, 0.45 μm pore size (Catalogue number: HABP04700; Millipore, Burlington, Massachusetts, United States). Explants were cultured in Dulbecco’s Modified Eagle’s Medium (DMEM; Sigma-Aldrich, Burlington, Massachusetts, United States) supplemented with 2% foetal bovine serum (FBS; Thermo Fisher Scientific, Waltham, Massachusetts, United States) and 1% penicillin-streptomycin (Thermo Fisher Scientific) at 37°C, 5% CO_2_.

For stretching experiments, skin was dissected and left free-floating in PBS for 20 minutes to permit full relaxation of the tissue. Skin explants were then manually stretched onto a nitrocellulose filter using fine-tipped forceps.

### mTmG cell labelling

To sparsely label cells with membrane GFP in mTmG mouse skin, explants from embryonic day **(E)** 13.5 were dissected as previously described, before being gently placed epidermis side down. The dermis was injected with approximately 2-5 μl TAT-Cre recombinase (200 U/ml; Sigma-Aldrich) using a glass microcapillary pipette. Explants were then attached to nitrocellulose filters epidermis side up and incubated for 24 hours at 37°C, 5% CO_2_ before being imaged.

### Chameleon cell labelling

Chameleon eggs were incubated until E3 before being windowed and the somites injected with approximately 2-5 μl TAT-Cre recombinase (200 U/ml) using a glass microcapillary pipette. Eggs were resealed and incubated at 37.8°C until E7 when skin explants were dissected and attached to nitrocellulose filters as described above.

### Immunohistochemistry

Samples were fixed overnight in 10% neutral buffered formalin, washed in PBST (PBS + 0.5% Triton x100) and incubated in blocking buffer (10% goat serum/PBST) for 2 hours at room temperature. Samples were incubated in anti-mouse collagen type I (AB765P; Sigma-Aldrich) in blocking buffer (1:50 dilution) overnight at 4°C, washed in PBST and incubated in goat antimouse IgG Alexa Fluor™ 647 (A21236; Thermo Fisher Scientific) in blocking buffer (1:250 dilution) overnight at 4°C. Samples were washed in PBST, counterstained with DAPI (1:2000; Sigma-Aldrich), washed in PBST and mounted on Superfrost slides (Thermo Fisher Scientific) with ProLong™ Gold Antifade mountant (Thermo Fisher Scientific).

### Imaging

Live time-lapse imaging sequences of TCF/Lef::H2B-GFP mouse and Chameleon chicken skin explants from Glover et al. and Ho et al. respectively ^6,11^ were used for tracking of dividing and non-dividing cells for Figures 1 and 2. Images were captured at 10 minute intervals for mouse time-lapse imaging and at 15 minute intervals for chicken.

For time-lapse imaging of mTmG explants and stretched TCF/Lef::H2B-GFP dorsal skin, explants on membrane filters were submerged in DMEM, supplemented with 2% FBS and 1% penicillin-streptomycin, in glass-bottomed 24-well plates (Eppendorf, Hamburg, Germany), epidermis down. Explants were imaged at 10 minute intervals using an LSM-880 confocal microscope (Zeiss, Oberkochen, Germany) in an incubation chamber at 37°C.

For whole embryo live time-lapse imaging experiments, embryos were harvested in PBS and affixed to a petri dish using superglue, before being submerged in DMEM, 2% FBS, 1% penicillin-streptomycin. Embryos were imaged at 5 minute intervals using an LSM 7 MP multiphoton microscope (Zeiss) with a perfusion system to maintain embryo temperature at 37^°^C (See Figure 3B).

The percentage decrease in mouse skin explant size after dissection was determined by imaging freshly dissected explants at 3 minute intervals using an Axiozoom V16 (Zeiss), and averaging 5 measurements across the anterior-posterior (A-P) axis and perpendicular to the A-P axis for each time point. The degree of stretch was obtained by imaging explants before and after stretching, averaging 5 measurements across the anterior-posterior (A-P) axis and perpendicular to the A-P axis each.

Fixed tissue samples were imaged using an LSM-880 confocal microscope.

### Cell tracking and division analyses

Cell tracking and division analysis was performed using custom written macros for ImageJ. Source code is available through GitHub (https://github.com/richiemort79/mitosis_tools). To analyse mitosis angles, maximum intensity Z-projections of time-lapse sequences were drift-corrected and all visible cell divisions were mapped by connecting the middle of daughter nuclei in the first frame of visualisation with a line. The angle of this line relative to x and y axes of the imaging frame was calculated using ImageJ.

To track cell behaviour, all visible dividing cells, their resulting daughter cells and at least 50 non-dividing cells were tracked manually until either the video ended, the cell entered a condensate or the cell migrated beyond the boundary of the video.

### Spatial analyses

Experimental angular variograms^28^ were produced on a point process represented by the mitosis locations in relation to the follicle, i.e. the variogram bins are calculated not between mitoses but between mitoses and follicle. Directional variograms are calculated considering the mitoses at four directions in respect to the follicle: 0, 45, 90 and 135 degrees, each of which within a cone of + and - 22.5 degrees. Experimental omnidirectional and directional variograms were estimated using the method of moments and fitted with exponential function by minimization of the weighted (by number of pairs in each bin) sum of squares^29^. From the fitted variograms the spatial range of influence of the follicle on the variance of the mitoses directions and the explained variance of the mitoses directions by follicle distance has been estimated. The latter is expressed in terms of nugget-to-sill ratio^30^.

Permutation analyses was employed to identify whether the modelled variogram (and its parameters) were statistically different, at a significance level of 0.001, from a random process^31^. Directions were permuted across all mitoses and for each permutation the experimental variogram was estimated and fitted. This process was repeated 1000 times. We estimated the Monte Carlo probability that permuted allocations of the directions produced smaller ranges, i.e. the probability that follicles were less influential on the mitoses direction. This is calculated as the number of permutations with lower spatial range divided by the total number of permutation (1000). If this value is lower than 0.001 then the null hypothesis (no influence) can be rejected.

Maps of the direction of the mitoses around the follicle (Figure 2D) were produced by conventional ordinary kriging.

### Local alignment analysis of *in-vivo* embryo mitoses

The random process of embryo mitotic directions, conditional to the mean and variance of the embryo mitoses, was estimated by permuting the directions and predicting them in a 100 by 100 pixel grid via conventional ordinary kriging. The latter uses the modelled variogram of the in-vivo embryo mitoses (unchanged). This process was repeated 100 times and the predicted average direction was estimated for each pixel^32^. Finally, we tested if the mean of the distribution of predicted mitosis directions derived from a random process in each pixel was different from the experimental mean of the direction of the mitoses in the same pixel^33^. The map of pixels containing directions that were not significantly different (at a 0.01 significance level) from a random process is shown in Figure 3F.

### Nuclei length and orientation analyses

Per skin sample, nucleus shape and orientation of the longest nucleus axis of 50 individual cells were measured using ImageJ. The length and angle of the cells were gathered and compared to the A-P axis direction.

### Alignment score analysis

Global alignment quantification for cell division angles and orientation of nuclei was conducted by analysing the vector movement of all nuclei within a dataset. Using all paired permutations of vectors and calculating the respective dot products, we computed an overall “alignment coefficient”:

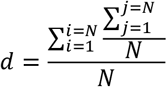

Where i and j = the upper and lower bounds, respectively, of the summation series, and N = total number of permutations of vector pairs. An alignment coefficient of 1 represents an ideally linear case, where all vectors are oriented across one primary angle. An alignment coefficient of 0 represents ideal isotropy where we observe a truly random distribution of vectors.

We used a colour-based segmentation approach to compute the overall alignment of stained collagen fibrils. Images were converted from the RGB colour space to the L*a*b colour space where we isolated the *a-channel* and negative region of the *b-channel*. Images were converted to a grayscale representation and passed through a median filter of order *n* = 2 to minimise the levels of salt and pepper noise. Contrast and brightness enhancement was applied with island size removal performed to maximise the isolation of fibrils from the background and nuclei. Images were converted to the frequency domain using a 2-dimentional Fourier transformation followed by transformation to polar coordinates. Using the polar form, the column intensities of all pixels were summed, representing the overall image intensity at each 1-degree bin size. The intensity values obtained relate to the proportion of collagen fibres which are oriented across each specific bin. These values were then used to calculate an overall alignment coefficient.

### Statistical methods

To compare the rate of condensate entry between newly born daughter and non-dividing cells, Fisher’s exact tests were performed between the frequency of daughter cells entering a condensate in the first 180 minutes post division, and the frequencies of non-dividing cells entering the condensate in 180 minute time windows throughout the imaging period. Fisher’s exact tests were also performed to assess timing of condensate entry after mitosis between the frequencies of dividing cells entering a condensate in the first 180 minutes post division and the frequencies of the same class of cells in later 180 minute time windows.

Unpaired Student’s T-tests were performed to determine p-values for average nucleus length values and alignment scores.

For comparing cell migration speed in stretched and relaxed skins, log cell speed was analysed using a mixed model with: frame, group, the frame x group interaction and log predivision speed (based on the mean of frames -4 to -1) fitted as fixed effects; and skin piece, family, cell ID, and the ‘frame x skin piece’ interaction fitted as random effects. A ‘family’ is made up of a parent cell its two daughters. Inclusion of the random effects allowed for additional random variation between skin pieces, family and cell IDs. In order to avoid the minus infinity values occurring for zero speeds, 0.01 was added to speeds before taking logs. The analysis was carried out using the MIXED procedure in the SAS/STAT software package, Version 9.4, Copyright © (2002-2012), SAS Institute Inc.

## Acknowledgements

We thank staff of the Biological Research Facility and Transgenic Chicken Facility in the National Avian Research Facility at the Roslin Institute for technical support, and Lorraine Rose for providing us with mTmG mouse embryos. We thank Raphael Voituriez for insightful discussion on cell movement.

This work was supported by BBSRC awards BB/T007788/1 and BBS/E/D/10002071, North West Cancer Research Fund award CR1132 and Royal Society, award RGS\R2\212427. SRN is supported by a scholarship from the EPSRC Centre for Doctoral Training in Statistical Applied Mathematics at Bath (SAMBa), under the project EP/S022945/1.480. This research made use of the Balena High Performance Computing (HPC) Service at the University of Bath. CAB is supported by a Carnegie Trust for Scotland PhD studentship.

