## Appendix for "Newly born mesenchymal cells disperse through a rapid mechanosensitive migration"

Affiliations:
[1] The Roslin Institute and Royal (Dick) School of Veterinary Studies, University of Edinburgh, Edinburgh, UK
[2] Department of Mathematical Sciences, University of Bath, Claverton Down, Bath, United Kingdom.
[3] Lancaster Ecology and Epidemiology Group, Lancaster Medical School, Lancaster University, United Kingdom.
[4] School of Engineering and Physical Sciences, Heriot-Watt University, Edinburgh, UK
[5] Division of Biomedical and Life Sciences, Faculty of Health and Medicine, Lancaster University, Lancaster, UK.

Hair follicle localisation involves the evolution of both temporal and spatial variables, such as cells' positions, the number of cells via division as well as the size of the condensates. It is important to model these variables to understand the process governing the localisation of cells to developing dermal condensates. Spatio-temporal mathematical models provide us a framework to model a variety of biological settings. Models where individual cells are viewed as agents on a domain are popular in mathematical biology. In this report, we adapt a spatiotemporal approach to model the process of hair follicle localisation and provide an estimation for all the modelling parameters. A table of parameters can be found on Page 8.

### 1 Model Description

We use an off-lattice agent-based method to model the localisation of the mesenchymal dermal cells to the developing hair follicles. The model has three physical components: (1) the developing dermis, (2) the hair follicles and (3) the moving cells each of which entails its own function and dynamics. We model the dermis as a square of fixed side length $L$ and the hair follicles as $M$ discs with a common radius $r_0$ whose locations on the domain are fixed. To simplify the modelling process, we assume that the follicles are present from the beginning and that they do not form or grow over time. We model the cells as points on the domain that do not exclude volume, are highly mobile and are able to divide. The initial positions of the cells are assigned uniformly at random. The boundary conditions on the boundaries of the domain are periodic and the boundary conditions on the discs representing the developing hair follicles are fully adsorbing, i.e., if a cell hits a developing hair follicle it is immediately adsorbed. When this happens, the adsorbed cell is considered 'removed' from the system and it loses its ability to move and divide.
For the off-lattice model, the coordinates of the cells' positions evolve in the continuous space following Langevin dynamics [1]. If $\left(X_i(t), Y_i(t)\right)$ denotes the position of the cell $i$ at time $t$, then the position of the cell at the time $t + \Delta t$ is given by,

$$X_i(t+\Delta t) = X_i(t) + \sqrt{2D\Delta t}\,\xi_x,$$
$$Y_i(t+\Delta t) = Y_i(t) + \sqrt{2D\Delta t}\,\xi_y,$$

where $\Delta t$ is the time step, $D$ is the diffusion coefficient and $\xi_x, \xi_y$ are independent and identically distributed Gaussian random variables with zero mean and unit variance.

At uniformly distributed random times, a cell is chosen to divide. For a dividing cell, three things happen sequentially in this order:

1. **Division:** The cell divides and the mother cell are replaced by two daughter cells.

2. **Mitotic jump:** One of the daughter cells travels a distance $d_{\text{mit}}$ in a straight line in a random direction with angle $\theta$ to the horizontal sampled from a given distribution peaked around $\pi/2$ rad (see Section 2.2) for a duration $\Delta t_{\text{mit}}$. At the same time the other daughter cell jumps the distance $d_{\text{mit}}$ in the diametrically opposite direction, $\theta + \pi$. We refer to this post-division movement as the 'mitotic jump' (Figure 1a). For a division event happening at time $t$, the position of the daughter cell $A$ and $B$ at the end of the jump is given by,

$$x_A(t + \Delta t_{\text{mit}}) = x(t) + d_{\text{mit}} \cos(\theta)\, \Delta t_{\text{mit}}, \quad y_A(t + \Delta t_{\text{mit}}) = y_A(t) + d_{\text{mit}} \sin(\theta)\, \Delta t_{\text{mit}},$$

   and,

$$x_B(t + \Delta t_{\text{mit}}) = x(t) + d_{\text{mit}} \cos(\theta + \pi)\, \Delta t_{\text{mit}}, y_B(t + \Delta t_{\text{mit}}) = y_B(t) + d_{\text{mit}} \sin(\theta + \pi)\, \Delta t_{\text{mit}},$$

   where $\big(x(t), y(t)\big)$ is the mother cell's position.

3. **Persistence:** Following the mitotic jump, each cell moves according to a persistent random walk. During the persistence phase, a cell moves in a given direction dictated by the angle of persistence, travelling in this direction for some amount of time, changing direction and moving in the new direction for some amount of time and changing direction again continuing with this motion for the total persistence time $T_{\text{pers}}$. The angle of persistence is defined as the amount of deviance from the division angle and is sampled from a distribution peaked around 0 rad (See Figure 6).

   If we let $\boldsymbol{\tau}_{\text{pr}} = \big(\Delta t_{\text{pers}}, 2\Delta t_{\text{pers}}, \dots, P\Delta t_{\text{pers}} = T_{pers}\big)'$ denote a uniform grid of the times at which a persisting cell changes direction and by $\boldsymbol{\phi} = (\phi_1, \dots, \phi_P)'$ the vector containing the angles of movement, then we can describe the persistent motion as follows. Immediately after the mitotic jump, the cell moves a distance $d_{\text{pers}}$ in a straight line in the direction defined by the angle $\phi_1$ moving in this direction for $\Delta t_{\text{pers}}$ amount of time, then changing direction and continuing moving in the new direction given by the angle $\phi_2$ and changing direction again at time $t = 2\Delta t_{\text{pers}}$ continuing with this persistent motion for a total time $T_{\text{pers}}$. A diagram showing a persisting cell is shown in Figure 1b. If we let $x_A(t)$ denote the position of daughter $A$ at the end of the mitotic jump, then the final position of the daughter cell at the end of the persistent period is given by,

$$x_A\big(t + P\Delta t_{\text{pers}}\big) = x_A(t) + d_{\text{pers}} \sum_{p=1}^{P} \cos\big(\phi_p\big)\, \Delta t_{\text{pers}},$$
$$y_A\big(t + P\Delta t_{\text{pers}}\big) = y_A(t) + d_{\text{pers}} \sum_{p=1}^{P} \sin\big(\phi_p\big)\, \Delta t_{\text{pers}},$$

   and likewise for the daughter cell $B$.

Not all cells on the domain have an equal chance of dividing. Cells that have been adsorbed or are already in any one of the three division phases described above cannot be chosen to divide. In this model a cell can divide more than once, and it is possible for the next division to occur immediately after the last division of the same cell. This is not biologically plausible as in reality a cell must progress through the cell cycle before dividing again. In this work, we have chosen the former description of cell division for simplicity in implementation of the model.

Biologically realistic estimates of the parameters of a model are needed to obtain plausible results and to give them meaningful interpretations. These estimates typically come from experimental data. While some parameters can be measured directly, such as the size of the domain, the number of cells and the radius of a hair follicle, others need to be estimated using an interplay of experimental data and mathematical and statistical methods. In the next section, we analyse experimental data to derive some of these parameters.

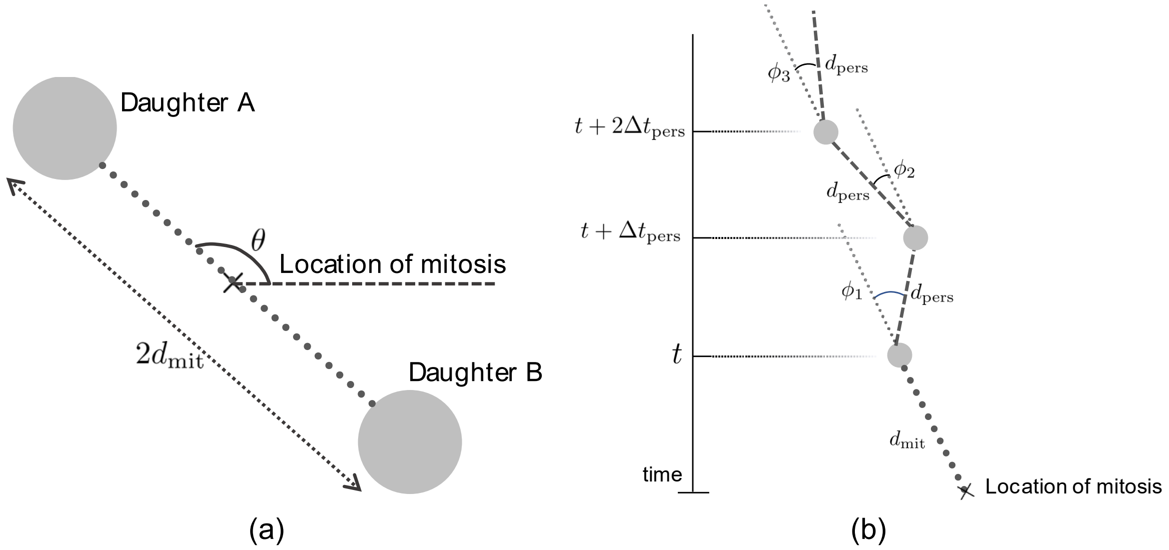


Figure 1: Two cells dividing and performing the mitotic jump (a) and a cell moving in a persistent way following the mitotic jump (b). In (a) cells A and B are separated by a distance $2d_{mit}$ at the end of the mitotic jump with each cell being a distance $d_{mit}$ away from the location of mitosis (marked with a black cross). Cell A moves in the direction given by the angle $\theta$ whereas cell B moves in the diametrically opposite direction, $\theta + \pi$. In (b), we show a possible trajectory (not drawn to scale) of cell A after division. After the mitotic jump (the grey dotted line) the cell undergoes a persistent random walk. The direction of travel is given by the angles $\phi_1, \phi_2$ and $\phi_3$ measured relative to the direction of the mitotic jump. The cell travels in this direction for an amount of time $\Delta t$ before changing direction again. The distance travelled before changing direction is $d_{pers}$.

### 2 Experimental Data Analysis

The ultimate goal for creating a mathematical model is to derive results reflecting the real biological process, in our case the localisation of mesenchymal dermal cells to the developing hair follicles. For the results to accurately reflect the real system, the model must be informed by experimental data. In this chapter, we apply well known mathematical techniques to the data to estimate our model parameters. In section 2.1 we use cell tracks to find an estimate for the diffusion coefficient $D$, in section 2.3 we characterise persistence of cells after division and in Section 2.2 we find the distribution of the division angles.

#### 2.1 Determining the diffusion coefficient

The diffusion coefficient $D$ quantifies how fast agents diffuse – a large value of $D$ corresponds to faster diffusion while a smaller value of $D$ corresponds to slower diffusion of agents. We estimate the diffusion coefficient by analysing cell tracks. The most widely used method for this is the mean square displacement (MSD) which tells us about the type of anomalous diffusion undergone by the cells [2] [3] [4] [5].

To calculate the MSD, different sections of the tracks of cells are considered using time windows of varying lengths. For a track consisting of $n$ successively recorded points,

$$r_k = (x_k, y_k), \quad \text{for } k = 1, \dots, n$$

the MSD for a window of length $\ell$ is given by,

$$\rho_\ell = \frac{1}{n-1} \sum_{k=1}^{n-1} (r_{k+\ell} - r_\ell)^2 . \quad \ell = 1, \dots, n-1$$

We fit a linear model of the form $\hat{\rho}(t) = bt$ to the calculated MSD where $\hat{\rho}(t)$ denotes a fitted value. The estimate of the diffusion coefficient is then recovered as $\widehat{D} = b/(2d)$ where $d$ denotes the dimensionality of the data, e.g., $d = 2$ for two-dimensional data.

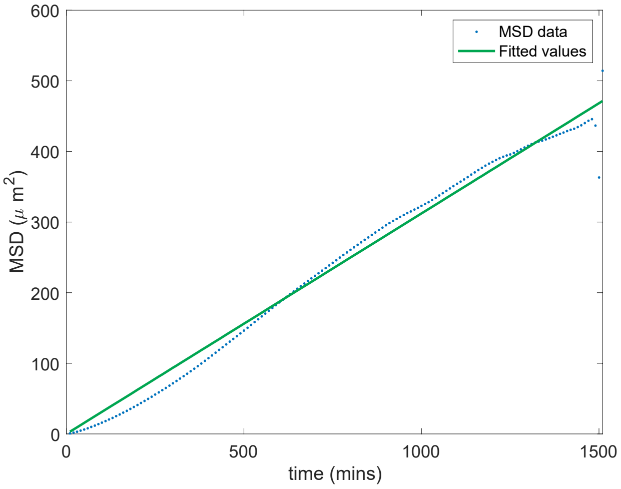


Figure 2: MSD of the 404 non-dividing cell tracks. The blue dots represent the calculated values, and the green line shows the fitted line with the equation $\hat{\rho} = 0.312t$.

Using the MSD, we analysed tracks of 404 cell non-dividing cells. The tracks were recording using a timestep of $10$ minutes before successive observations. The number of data points in each track was variable with the smallest track comprising only 3 pairs of $x$ and $y$ coordinates and the longest track having 152 pairs. The majority of the tracks were large. Figure 2 shows the ensemble-averaged MSD (blue dots) with the fit given by the equation $\hat{\rho}(t) = 0.312t$ (solid green line). We find the average diffusion coefficient to be 0.0780 $\mu$m$^2$min$^{-1}$ (4 d.p.) with a $95\%$ confidence interval [0.0773,0.0788].

#### 2.2 Division

Here we find the necessary parameters i.e., the division angles $\theta$ and the size of the mitotic jump $d_{\text{mit}}$ to delineate division and the mitotic jump. Using the last position of the mother cells immediately prior to dividing and the first position of the daughter cells' tracks post-division we were able to calculate the size of the mitotic jump for 662 daughter cell tracks. The mean jump size was found to be 6.4271 $\mu$m.

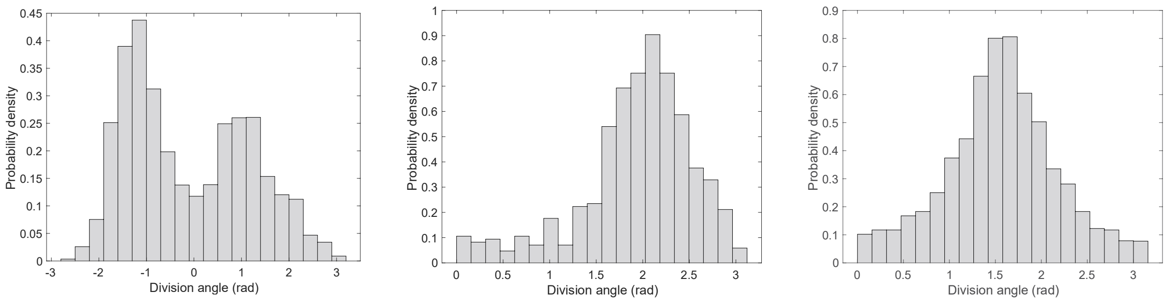


(a) (b) (c)

Figure 3: Shifting and transforming division angles: (a) raw angles $\Theta_0$ with distribution $\mathcal{D}_0$, (b) angles after being shifted $\Theta_1 = (\Theta_0 + \pi) \bmod \pi$ with distribution $\mathcal{D}_1$ and (c) the final adjusted angles $\Theta_2 = (\Theta_1 + \pi/2 - \theta_m) \bmod \pi$ with distribution $\mathcal{D}_2$.

Figure 3a shows a histogram of division angles from 3,114 divisions. The histogram appears to be bimodal with one peak at $-1.19$ rad ($-68.24$ degrees) and another one at 0.99 rad (56.93 degrees). One reason for this bimodality is that upon division the pair of daughter cells are spawned in diametrically opposite directions meaning their angles of motion are a phase difference of $\pi$ rad (180 degrees) apart. Often measuring the angle for the cell spawned in one direction while measuring some angles for cell spawned in the opposite direction could result in a distribution with two peaks. Secondly, a shift in the histogram can occur due to the way images of the specimen were captured. In most cases, the images were taken to align the division angles about the vertical axis of $\pi/2$ rad (90 degrees). However, in some situations, a different orientation of the images meant the angles were misaligned resulting in a shift in the angles from these images.

To treat this bimodality, we apply a series of transformations to the set of unadjusted and raw angles $\Theta_0$. To discard one of the peaks we first apply the transformation $(\Theta_0 + \pi) \bmod \pi$ which shifts and merges the angles that were a phase difference apart. The distribution, $\mathcal{D}_1$ for the updated set of angles$\Theta_1$, is shown in Figure 3b. This is followed by applying the transformation $(\Theta_1 + \pi/2 - \theta_m) \bmod \pi$, where $\theta_m$ is the angle at the maximum of $\mathcal{D}_1$. This transformation aligns all the angles about a common axis of $\pi/2$ rad (90 degrees), yielding the updated set of angles, $\Theta_2$ with the distribution $\mathcal{D}_2$ whose circular mean $\mu$ is given by 1.56 rad (89.52 degrees) (Figure 3c).

To sample angles from this distribution, we need to fit an appropriate probability density function (pdf) to the observed distribution. Circular distributions, also known as wrapped distributions, are a group of distributions used to model directional data. There are many such distributions such as the cardioid distribution, the wrapped Cauchy distribution, the von Mises distribution and the wrapped Gaussian distribution. For this work, we choose the von Mises distribution which is used widely for data involving angles [6].

In this context, the $\mu$ is simply the circular mean of the division angles which is $1.562$ rad ($87.4$ degrees). To find $\kappa$ we seek a value that would minimise the sum of squared residuals (SSR) between the fitted distribution for the different possible values of $\kappa$, $\boldsymbol{k} = 0{:}0.01{:}3$ and the empirical distribution,

$$SSR = \sum_{i=1}^{m} \left( \hat{y}(\phi_i; \mu, k_j) - y(\phi_i)^2 \right), \qquad j = 1, \dots, len(\boldsymbol{k})$$

where $\hat{y}(\phi_i; \mu, k_j)$ is the value of von Mises probability density function (pdf) at the angle $\phi_i$ evaluated for $\mu$ and $k_j$ and $y(\phi_i)$ is the value of the observed distribution at the angle $\phi_i$.

Using the above method we find that $\kappa^\star = 2.85$ provides the most appropriate fit. The optimal fit is shown in Figure 4 overlaid on the histogram for comparison.

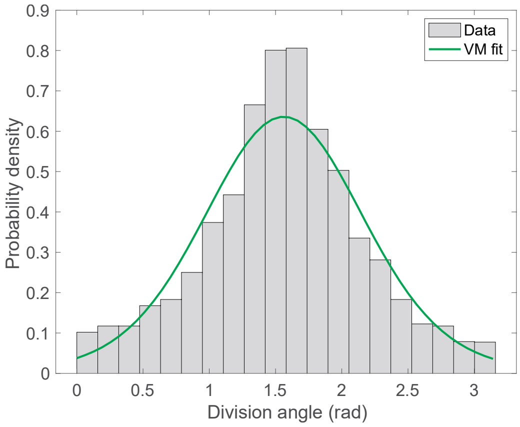


Figure 4: Distribution of division angles after adjustment. The histogram depicts the empirical distribution whereas the curve represents the von Mises distribution with $\mu = 1.562$ rad (89.52 degrees) and $\kappa = 2.85$.

#### 2.3 Characterising persistence

The MSD can be used to determine the type of anomalous diffusion undergone by the cells. Generally, the relationship between the MSD and time can be written in the form $\rho(t) = bt^\alpha$ where $b > 0$ and $0 \leq \alpha \leq 2$. In the case of standard diffusion, $\alpha = 1$ and the MSD curve is linear taking the form $\rho(t) = bt$ as we have seen in Section 2.1. However, in situations in which the dispersal of cells is anomalous, the MSD will not follow a linear relationship. Broadly speaking, anomalous diffusion can be divided into two subcategories: sub-diffusion and super-diffusion [4] [5]. Sub-diffusion occurs when the MSD is increasing at a slower rate than standard diffusion (i.e.$0 \leq \alpha < 1$) whereas in super-diffusion the MSD is increasing at a faster rate than standard diffusion (i.e.$1 < \alpha \leq 2$). Persistent motion is an example of super-diffusive behaviour.

We calculate the MSD for the dividing cells utilising the 180 minutes after the mitotic jump. If a track is shorter than 180 minutes, we simply calculate up to the maximum length of the track available. In order to determine if the cells are persisting, we fit a linear model of the form $\log(\rho) = \alpha \log(t) + \log(b)$ to the experimentally calculated MSD. We find $\alpha = 1.5380$ (4 d.p.) and $b = 0.0853$ (4 d.p.). Since $\alpha > 1$, it is evident that the cells follow a persistent random walk following a mitosis. Figure 5 shows the MSD (blue dots) with the fitted curve (green curve). We can see that the curve is a good fit for the early part of the MSD data however the fit deviates from the data for larger times, indicating that persistence diminishes with time.

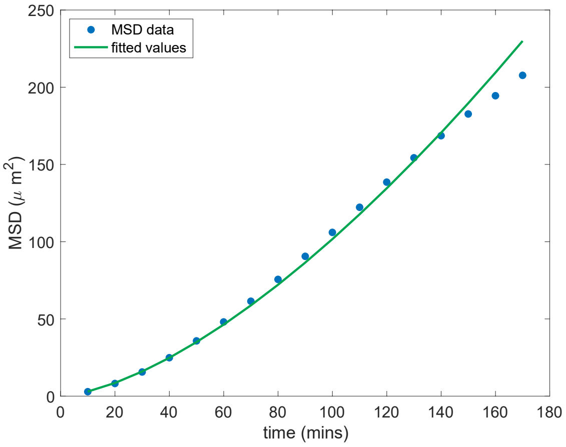

Figure 5: MSD of the 662 cell tracks for the first 180 minutes after the mitotic jump. The blue dots represent the data and the green curve is the fit with equation$\hat{\rho} = 0.0853t^{1.538}$.

Two key parameters are required to model the persistent motion of cells: the movement angle, $\phi$, relative to the angle of division and the distance moved before changing direction, $d_{\text{pers}}$. Knowledge of those two parameters is necessary to fully delineate persistence.

#### 2.4 Persistence Angle

By analysing cell track data after the mitotic jump for the next 180 minutes, we were able to find the distribution of the persistence angle $\phi$. The persistence angle $\phi$ is defined as the derivation in the direction of a track from the direction of its mitotic jump. Figure 6 shows a histogram of these angles over 662 tracks. The distribution suggests that the direction of motion of persistent cells is centred around approximately 0 rad with the mean and variance taking values $-0.078$ rad (3 d.p.) and 0.612 rad (3 d.p.), respectively. The centring of the distribution of these angles about an angle close to 0 suggests that for the next 180 minutes after dividing, the cells do not deviate much from the direction of their mitotic jump.

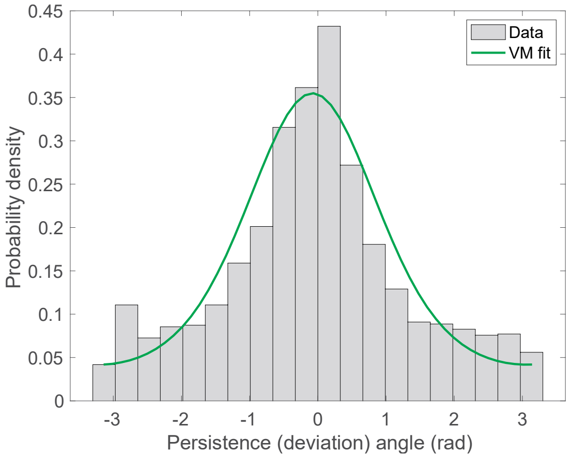


Figure 6: Distribution of persistence angles. The histogram depicts the distribution of persistent angles whereas the curve (in green) is the VM fit with $\mu = -0.0782$ rad ($-4.48$ degrees) and $\kappa = 1.07$.

Using the method highlighted previously in Section 2.2, we fit a von Mises distribution. We found $\mu = -0.0782$ rad and $\kappa$ = 1.07 to provide an optimal fit. In Figure 6 we show the histogram of the persistence angles and we overlay the optimal von Mises pdf for comparison. The VM distribution seems to provide a good fit by capturing the important features of the empirical distribution.

#### 2.5 Distance

We used the persistent tracks to estimate the average distance moved by a cell in a given direction before changing its direction. The distance between each pair of successively recorded position was calculated over all the 662 tracks. We find the mean distance to be 1.5384 $\mu$m.

In the next chapter we present and discuss the results obtained via the implementation of the model with the parameters derived here. A full list of parameters is given in Table 1.

Table 1: Table of all model parameters

| Parameter description | Parameter symbol | Parameter value |
|---|---|---|
| Domain side length | $L$ | $634.15\ \mu$m |
| Number of follicles | $M$ | 14 |
| Radius of a follicle | $r_0$ | 35 microns |
| Initial number of cells | $N_{init}$ | 1936 |
| Diffusion coefficient | $D$ | 0.078036 microns$^2$min$^{-1}$ |
| Angle of division | $\theta$ | $VM(1.5625, 2.8500)$ |
| Size of mitotic jump | $d_{\text{mit}}$ | 6.4271 microns |
| Length of mitotic jump period | $\Delta t_{\text{mit}}$ | 10 mins |
| Total modelling time | $T_{final}$ | 1800 minutes |
| Angle of persistence | $\phi$ | $VM(-0.0782, 1.0700)$ |
| Length of time before changing direction during persistent motion | $\Delta t_{\text{pers}}$ | 10 minutes |
| Distance covered during a persistence period | $d_{\text{pers}}$ | 1.5384 microns |
| Total persistence time of a cell | $T_{\text{pers}}$ | 180 minutes |

### 3 Results

We implement our model in the high-level programming language MATLAB in order to answer the biological questions surrounding the localisation of cell to dermal condensates.

Experimental data suggests that dividing cells are more likely to be adsorbed by a dermal condensate than the non-dividing cells. The observed proportion of adsorbed dividing cells to adsorbed non-dividing cells is 19.37% to 14.30% (see Figure 7). We are interested in exploring the mechanism responsible for this disparity. One hypothesis suggests that the mitotic jumps are responsible for the higher proportion of dividing cells. In the next section, we use our mathematical model to test this assertion. Furthermore, to learn how the initial positions of cells affects the likelihood of adsorption, we use simulated data to find the distribution of the initial location of cells conditional on the fact that they are eventually adsorbed.

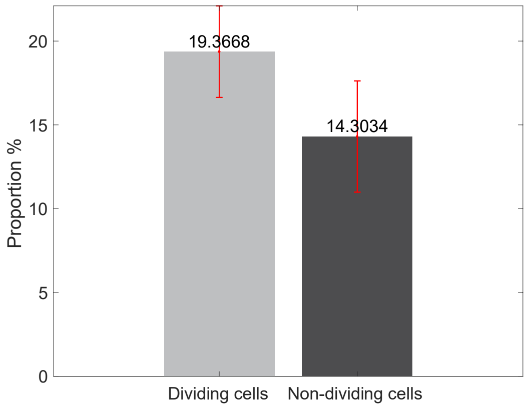

Figure 7: The proportions of dividing and non-dividing cells entering dermal condensates calculated using data from 8 experiments all performed under the same conditions. The error bars show mean $\pm$ standard error.

To compare our results with the experimentally observed proportions, we calculate the percentage of dividing and non-dividing cells that eventually get adsorbed by a developing hair follicle. To do this, we sample the same number of dividing cells as non-dividing cells, and we track their movement over time. Define $N_{\mathrm{pa}}$ as the number of cells that proliferated and got adsorbed, $N_{\mathrm{npa}}$ as the number of cells that did not divide and got adsorbed and $N_{\mathrm{mit}}$ as the total number of dividing cells being tracked. Then,

$$P_{\mathrm{pa}} = \frac{N_{\mathrm{pa}}}{N_{\mathrm{mit}}},$$

is the proportion of dividing cells that got adsorbed and,

$$P_{\mathrm{npa}} = \frac{N_{\mathrm{npa}}}{N_{\mathrm{mit}}},$$

is the proportion to non-dividing cells that got adsorbed.

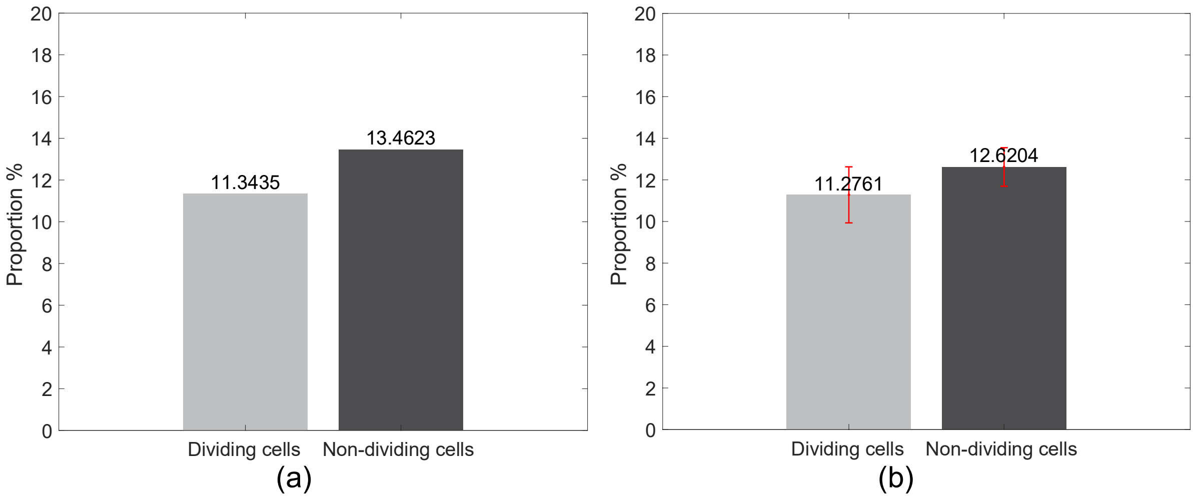


Figure 8: Proportions of adsorbed dividing and non-dividing cells for simulated without persistence from 1,000 repeats (a) and proportions averaged over 8 randomly chosen repeats with the corresponding 95% confidence interval (b).

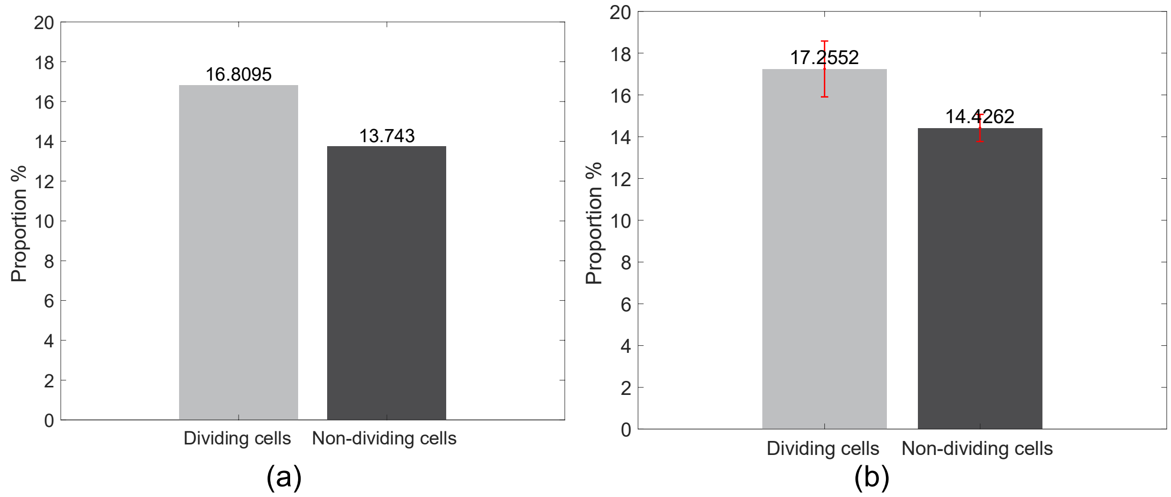


Figure 9: Proportions of adsorbed dividing and non-dividing cells with persistence following a mitotic jump averaged over 1,000 repeats (a) and over 8 repeats with 95% confidence interval (b). In both plots, we see a good representation of the observed proportions.

To assess the effect of the mitotic jumps on the proportions of cells adsorbed, we simulate the model without persistence. i.e., cells would diffuse and divide whereupon the two daughter cells would perform a mitotic jump but thereafter would not display the persistent movement associate with a jump. The calculated proportions are shown in Figure 8. We show proportions averaged over 1,000 repeats for greater certainty but also for a much smaller number of repeats to show how the uncertainty in the model results compares with the observed proportions. We immediately notice that the simulated proportions do not reflect the observed proportions (Figure 7). According to the simulated proportions, the dividing cells with mitotic jumps are less likely to be adsorbed than their non-dividing counterpart. This contradicts the hypothesis that the division jumps are responsible for a higher proportion of dividing cells.

To discern what role persistence plays in the hair follicle localisation process we introduce persistence into the model and calculate the mean percentage of adsorbed dividers and non-dividers. The results are presented in Figure 9. The proportion of adsorbed dividing cells is now higher than the proportion of non-dividing cells and is a good representation of the observed proportions.

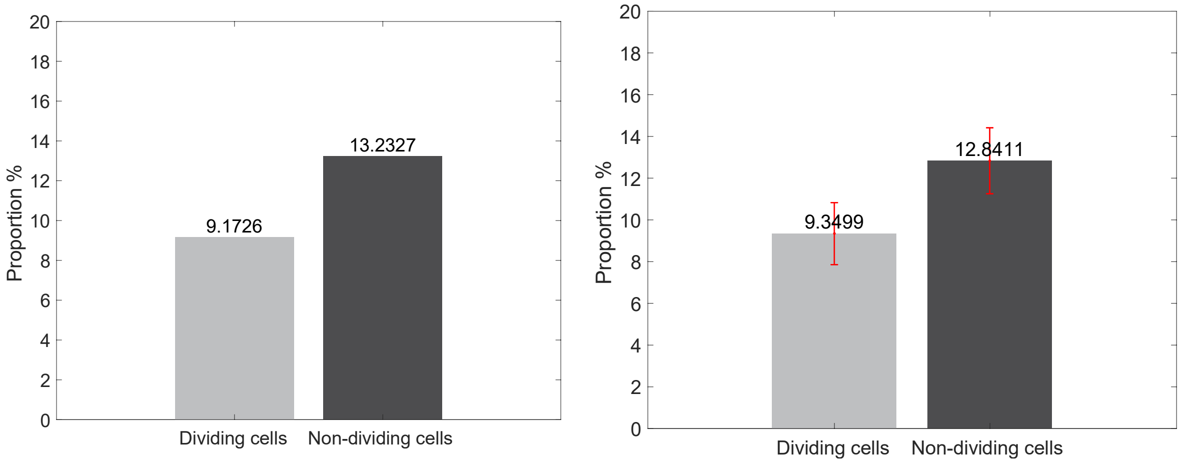


(a) (b)

Figure 10: Proportions of adsorbed dividing and non-dividing cells with no daughter cells produced at division averaged over 1,000 repeats (a) and over 8 repeats with 95% confidence bounds (b).

Considering a hypothetical scenario where no new cells are created at division and the mother cell performs a solo mitotic jump and persists, we see a much lower proportion of dividing cells compared to their non-dividing counterpart (see Figure 10). Hence, we conclude that mitotic jumps and persistence is not enough for the adsorption of cells. The higher proportion of dividers we see in Figure 9 is also due to having more dividing cells.

#### 3.1 Initial positions of cells

To explore how the initial position of cells affect their chances of adsorption, we find the distance between the initial location of a cell and its nearest condensate. To this end, we first calculate the distance between the initial location of each cell and the centre of each follicle for cells that got adsorbed at a time greater than 0. Then, for each cell we take the smallest of these distances to gives us the initial distance between the cell and its nearest dermal condensate.

To calculate the distribution of the initial distances given adsorption, we bin these distances into annuli of width $dr$ where the radius of each annulus is given by $r_0 + p \times dr$, where $r_0$ is the radius of the follicle and $p = 1, \dots, P$ with $P$ being the number of annuli around the condensate (Figure 11a). The bin counts are then normalised by the area of each annulus. The distribution itself is normalised by the total area under the resulting curve.

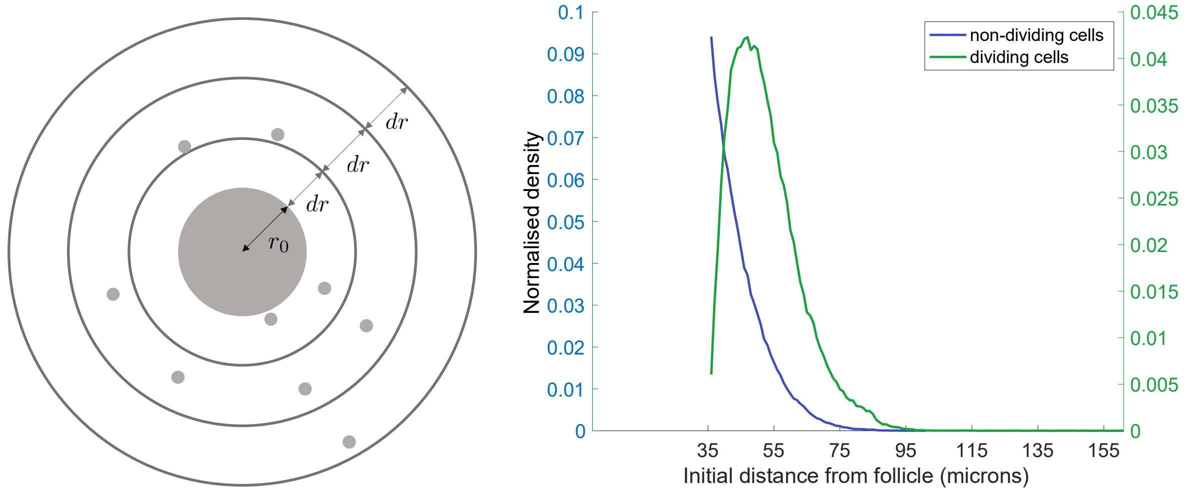

(a) (b)

Figure 11: (a) Binning the initial distances of cells. The big grey circle in the centre with radius $r_0$ is the dermal condensate and the smaller grey dots are the mesenchymal cells. The concentric rings are the annuli and the width of each annulus is $dr$. (b) Distribution of initial distance from the nearest hair follicle give that the cell is then adsorbed for non-dividing cells and dividing cells over 1,000 repeats.

In Figure 11b we display the normalised distribution of the initial distances from the nearest condensate of adsorbed non-dividing and dividing cells. The distribution for non-dividing cells appears to be decreasing monotonically which suggests that cells very close to a condensate are more likely to be recruited than cells far away from it. Comparing this to the dividing cells, the distribution is non-peaked at around 50 microns suggesting adsorption is most likely at around this distance.

This behaviour can be explained using the distribution of the time to adsorption of cells. In Figure 12 we can see that adsorption mostly occurs at the start hinting that cells very close to a condensate are recruited more sooner, leaving little chance for these cells to divide before adsorption.

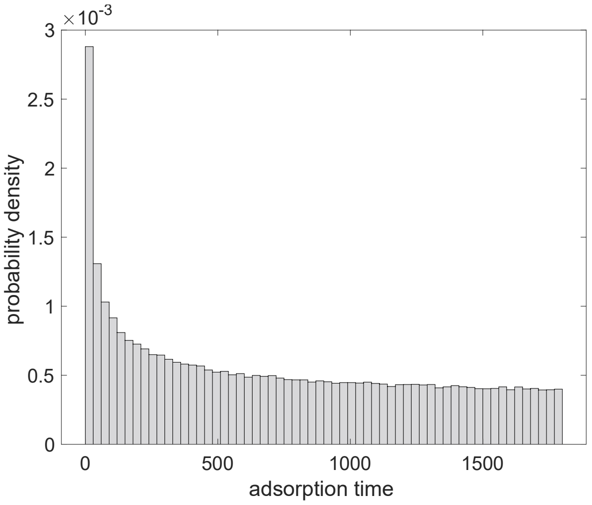


Figure 12: Distribution of time to adsorption of all the cells with non-zero adsorption time over 1,000 repeats.
